# Longitudinal assessment of the conversion of mild cognitive impairment into Alzheimer’s dementia: Observations and mechanisms from neuropsychological testing and electrophysiology

**DOI:** 10.1101/2024.12.16.628666

**Authors:** Dominic M Dunstan, Edoardo Barvas, Susanna Guttmann, Roberto Frusciante, Beatrice Viti, Mirco Volpini, Milena Cannuccia, Chiara Monaldini, Francesco Tamagnini, Marc Goodfellow, Luke Tait

## Abstract

**INTRODUCTION:** Elucidating and better understanding functional biomarkers of Alzheimer’s disease (AD) is crucial. By analysing a detailed longitudinal dataset, this study aimed to create a model-based toolset to characterise and understand the conversion of mild cognitive impairment (MCI) to AD.

**METHODS:** EEG, MRI, and neuropsychological data were collected from participants in San Marino: AD (n = 10), MCI (n = 20), and controls (n = 11). Across two additional years, MCI participants were classified as converters or non-converters.

**RESULTS:** We identified the Stroop Color and Word Test as the largest differentiator for MCI conversion (ROC AUC = 0.795). This was underpinned by disconnectivity in working memory and attention networks. Unsupervised clustering of EEG spectra also differentiated MCI conversion (ROC AUC = 0.710) and was underpinned by reduced excitatory and enhanced inhibitory synaptic efficacy in (prodromal) AD. Combining electrophysiological and neuropsychological assessments increased the accuracy of the differentiation (ROC AUC = 0.880) in comparison to each measure considered individually.

**CONCLUSION:** Combining electrophysiological and neuropsychological assessment with mathematical models can inform the development of non-invasive, low-cost tools for the early diagnosis of AD.

**Highlights:**

- We analysed longitudinal changes in EEG and neuropsychological assessments in MCI
- Stroop Color and Word Test error scores were lower in MCI converters
- The degree of impairment was found to be correlated with functional disconnectivity
- Unsupervised clustering of EEG spectra characterised patterns associated with disease
- Mathematical modelling revealed reduced excitatory synaptic efficacy in (prodromal) AD

**Research in Context:** **Systematic review:** The authors used PubMed to review the literature on the use of inexpensive modalities, including EEG and neurophysiological testing, for characterising the progression of MCI to AD. Although promising, existing work suggests the full potential of these methods as tools for understanding prodromal AD is still lacking.

**Interpretation:** A novel application of a clustering algorithm to EEG spectra revealed different patient diagnoses could largely be characterised by their cluster assignment. We also found differences in a particular neuropsychological test, the Stroop Color and Word Test. Using mathematical modelling we found there were both network and synaptic mechanisms that underlie these differences.

**Future directions:** Using the methods described herein to build markers for testing MCI to AD conversion on a large independent cohort will be crucial to understanding the full impact and applicability of these approaches. This may ultimately lead to a better characterisation and understanding of the diagnosis and prognosis of AD.

## 1 Introduction

Alzheimer’s disease (AD) is the most prevalent cause of dementia globally, yet it remains incurable. Recently approved disease-modifying therapies, such as Lecanemab (van Dyck et al., 2023) and Donanemab (Mintun et al., 2021; Sims et al., 2023), target the clearance of amyloid beta (Ab) peptides and plaques. However, their efficacy is limited, stage-dependent, and costly (Ross et al., 2022). Early intervention is crucial, as the initial stages of AD often precede clinical diagnosis by several years (Vermunt et al., 2019). Thus, we urgently need to better understand the progression of the disease in its early stages.

A definitive diagnosis of AD is currently possible only post-mortem by identifying extracellular Ab plaques and intracellular tau protein tangles in individuals with severe cognitive impairment at death (Hyman et al., 2012). Probable AD diagnosis can be established by combining clinical assessments, such as neuropsychological scores (NPS), with biological criteria and biomarkers obtained from cerebrospinal fluid (CSF) or positron emission tomography (PET) (Dubois et al., 2021). However, these two latter methods are either invasive or expensive, making them unsuitable for large-scale screening. Similarly, existing methods for identifying prodromal AD are not feasible for widespread screening due to their high cost and/or invasive nature. Additionally, while AD is the most common cause of dementia, it is not the only one, and mild cognitive impairment (MCI) does not always progress into dementia, with the median conversion of MCI to AD across two years at only 18.6% (Ward et al., 2013).

There is therefore a critical need to identify functional and biological markers for early-stage AD using non-invasive, low-cost techniques. This would aid in early diagnosis, differentiate MCI due to prodromal AD from other causes, and maximise treatment effectiveness. Understanding the mechanistic relationships between impaired brain function and AD-associated neurophysiological impairments and cognitive symptoms is also essential for developing better treatments. Electroencephalography (EEG) is a relatively inexpensive and non-invasive tool that has shown promise in early AD diagnosis (Babiloni et al., 2016; Cassani et al., 2018; Tait et al., 2020). EEG records electrophysiological oscillations of the brain via electrodes on the scalp. These oscillations, resulting from coordinated neuronal activity, can be analysed to understand different cognitive and perceptual states (Zani et al., 2020; Adamantidis et al., 2019). EEG is already used in neurology clinics for diagnosing conditions like epilepsy and multiple sclerosis. Its high temporal resolution and correlation with mental states make it a valuable tool for further investigation. In addition, EEG changes can capture neuronal degeneration caused by AD long before behavioural symptoms appear (Cassani et al., 2018). It has been consistently observed that EEG changes in (prodormal) AD patients include reduced power at high (alpha to gamma) frequencies and increased power at lower (delta to theta) frequencies (Pucci et al. (1998); Benwell et al. (2020); Babiloni et al. (2004); Bennys et al. (2001); Tait et al. (2019); Kopcanova et al. (2024)). These changes are thought to be associated with altered synaptic connectivity due to Ab’s deposition and neurofibrillary tangles (Choi et al., 2023; Tamagnini et al., 2015; Palop and Mucke, 2010; Gallego-Rudolf et al., 2024). Mathematical models have shown to be useful for interpreting synaptic and cellular mechanisms underpinning altered electrophysiological dynamics in neurodegenerative disease (Shaw et al. (2021); Ranasinghe et al. (2022); Bhattacharya et al. (2011)). Furthermore, when combined with neuropsychological testing, they allow the mechanisms underlying cognitive deficits to be inferred (Adams et al., 2021). Therefore, combining EEG, NPS and mathematical modelling could offer a comprehensive approach to understanding early AD progression and improve diagnosis in a cost-effective, non-invasive way that would be suitable for large-scale screening (Moretti et al. (2011); Rossini et al. (2006); Rossini et al. (2022)).

In this study, we aimed to characterise EEG and NPS changes in early AD and use mathematical models to understand any observed changes. We followed a sample population with MCI over two years, collecting EEG, NPS, and MRI data at multiple time points. We found that behavioural and electrophysiological measures can effectively track cognitive decline in individuals with MCI, and show how mathematical modelling can provide novel insights into the underlying mechanisms driving the observed functional changes.

## 2 Methods

### 2.1 Participants and recruitment

People with AD (n=10), amnestic MCI (n=20) and a control group of cognitively healthy older adults (HOA, n=11) were recruited from the Neurology Unit of the San Marino State Hospital. AD was diagnosed according to the DSM-5 criteria for major neurocognitive disorder due to Alzheimer’s dementia (American Psychiatric Association, 2013), and in-vivo evidence of Alzheimer’s pathology was also required. Specifically, inclusion criteria were: (1) a diagnosis of major neurocognitive disorder due to probable AD according to the DSM-5 criteria and (2) increased tracer retention on amyloid PET or a CSF Tau/A*β*_1–42_ ratio *>* 0.52. All of the people in the AD group had a diagnosis of typical AD according to the Dubois et al. (2014) IWG-2 criteria, they all showed clear evidence of memory decline and at least one other cognitive domain, a steady and gradual decline in cognition without extended plateaus, and no evidence of mixed aetiology. Nine patients had a CSF Tau/A*β*_1–42_ ratio *>* 0.52, which is considered a robust profile for AD (Duits et al., 2014). One patient had increased tracer retention on amyloid PET with 18F-florbetapir.

Inclusion criteria for people in the MCI group were: (1) cognitive changes fulfilling the criteria for mild neurocognitive disorder due to possible AD according to the DSM-5 criteria and (2) a performance that falls more than 1.5 standard deviation below age-appropriate norms in at least one memory test. For 4 MCI participants, CSF markers were also available, showing evidence of an Alzheimer’s pathology.

Finally, the HOAs in the study were recruited from caregivers of patients. Exclusion criteria for the HOAs were: (1) the presence of significant hearing or vision loss or (2) history or evidence of other neurological disorders or potential different causes for neurocognitive disorder or (3) an MMSE (Folstein et al., 1975) score below the normal limits (Magni et al., 1996).

After an extensive battery of neuropsychological testing (see subsection 2.2 for details), all the diagnoses were classified by a neurologist expert in neurodegenerative diseases. All AD patients were being administered anticholinesterase drugs in accordance with the Good Clinical Practice regulations of the Republic of San Marino.

After the initial baseline assessment (year 0), people with MCI were longitudinally followed at a one and two-year time point. These follow-ups entailed neuropsychological and clinical assessments to track the progression of MCI and test whether the DSM-5 criteria for major neurocognitive disorder due to probable AD had been met. People with MCI who converted to AD within two years are referred to herein as converters (MCI_c_) and people who did not are referred to as non-converters (MCI_nc_). Figure 1 illustrates the longitudinal data acquisition of the study. Participants provided written informed consent before participating and were free to withdraw at any time. All procedures were approved by the Republic of San Marino Ethical Committee for Research and Experimentation (Ref. 0015 SM) and the University of Exeter Medical School Research Ethics Committee.

**Figure 1:**
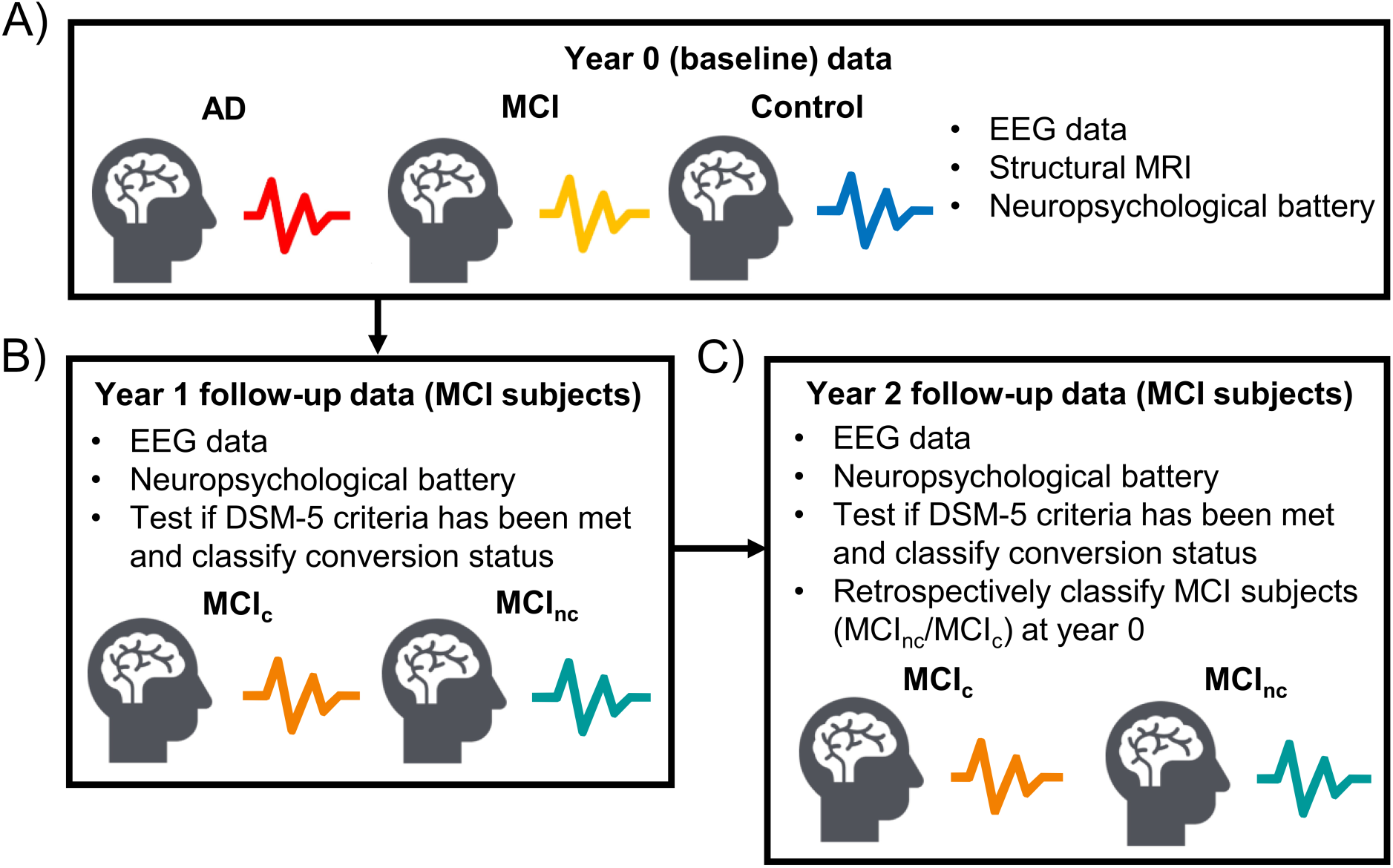
Schematic of longitudinal data acquisition. A-C) Detail the data obtained in the study across years 0, 1 and 2 respectively.

### 2.2 Neuropsychological assessment

Patients (MCI and AD) underwent a detailed neuropsychological battery. The battery included measures of verbal short-term memory (Digit Span Forward, DSF; Monaco et al. (2015)), working memory (Digit Span Backward, DSB; Monaco et al. (2015)), visuo-spatial short-term memory (Corsi Block-Tapping test, CBT; Monaco et al. (2015)), verbal learning (Rey Auditory-Verbal Learning Test – Immediate Recall, RAVLT-IR; G A Carlesimo (1996)), long-term verbal memory (Rey Auditory-Verbal Learning Test – Delayed Recall, RAVLT-DR; G A Carlesimo (1996)) and long-term visual memory (Modified Taylor Complex Figure Test – Delayed Recall, MTCF-DR; Casarotti et al. (2014)). Moreover, according to the procedure described by Ricci et al. (2012), after the RAVLT-DR the recognition trial was administered, and the Memory Efficiency Index (MEI) was calculated. Assessment of attention and executive functions included the Multiple Features Target Cancellation (MFTC; Marra et al. (2013)) scores for, correct hits (MFTC-H), errors (MFTC-E), execution time (MFTC-T) and accuracy (MFTC-A), the Stroop Color and Word Test (SCWT; Caffarra et al. (2002)) for errors (SCWT-E) and execution time (SCWT-T), and the Weigl’s Sorting Test (Weigl’s; Laiacona et al. (2017)). Language and praxis were assessed by means of a Pictures Naming test (PN; Catricalà et al. (2013)) and Ideomotor Apraxia Test (IMA; De Renzi et al. (1980)), respectively. Finally, visuo-constructive abilities were assessed using a copy of the Modified Taylor Complex Figure (MTCF-C; Casarotti et al. (2014)). Of these measures, HOAs only underwent MFTC and RAVLT, and an additional MMSE (Folstein et al. (1975)), to confirm normal cognitive functioning.

### 2.3 EEG recordings

EEG was recorded at baseline from all 41 participants using the EBNeuro BE Plus PRO device. EEG was recorded during rest, and participants were periodically instructed to either open or close their eyes. The mean duration of recording was approximately 17 minutes. For each participant, the EEG data was recorded at 256 Hz across 19 channels (Fp1, Fp2, F7, F3, Fz, F4, F8, T3, C3, Cz, C4, T4, T5, P3, Pz, P4, T6, O1, O2) according to the international 10-20 system. Recordings were viewed on a Galileo NT Line version 4.40/00/c31416 and then exported into MATLAB (MATLAB, 2024) for offline analysis. All epochs of artifact-free eyes-closed resting-state with a duration greater than 30 seconds were taken forward for analysis. Using a 4^th^-order Butterworth filter, the data was bandpass filtered between 1-70 Hz, notch filtered at 50 Hz and re-referenced to the common average. Independent component analysis was applied for artefact removal (including muscle and eye movements). EEG data analysis was performed using the Fieldtrip toolbox (Oostenveld et al. (2011), http://fieldtriptoolbox.org). To assess changes in EEG signals over time for the MCI group, people with MCI additionally underwent EEG recording using the same paradigm during their 1 and 2-year follow-up assessments. EEG data from the follow-up assessments were processed using the same protocol described above.

### 2.4 Power spectral analysis

Power spectral density (PSD) was estimated across a frequency range of 2-30 Hz for each participant using Welch’s method (Welch, 1967), across 30 s intervals. Within each interval, the signal was split into 8 segments with 50% overlap. A 0.25 Hz frequency resolution was used. The PSD was calculated for all 19 electrodes, normalised to unit total power, and then averaged (across intervals and channels) in the frequency domain to obtain a mean PSD for each participant.

To compare PSDs, the distance (denoted *D*) between the PSDs of participants *i* and *j* was calculated as

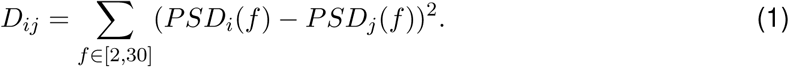

Note, this distance is commutative (i.e. *D_ij_* = *D_ji_*). To visualise and interpret differences between spectra, classical multi-dimensional scaling was used to embed *D* into two dimensions, whilst preserving the pairwise distance between subjects in the original space (Kruskal, 1964). Each participant was then represented by a scatter point in two dimensions, with the distances between points (participants) representative of the differences between the participants’ corresponding PSD.

We then used a Gaussian mixture model (GMM) to cluster the subjects in the two-dimensional space. To assess the optimal number of clusters, in addition to a visual inspection, we evaluated the Akaike information criterion (AIC) and the Bayesian information criterion (BIC; McLachlan and Rathnayake (2014)). These metrics estimate the predictive error of the statistical model, enabling the number of clusters to be determined by minimising this error. Based on the AIC and BIC, in this work, we choose to use *k* = 3 for the number of clusters (see Supplemental Figure S9 for more information).

### 2.5 MRI Acquisition and EEG Source Localisation

Participants underwent structural MRIs acquired with a GE Brivo MR355 scanner using a 1.5T magnetic field strength, using a T1-weighted axial sequence (echo time: 1.2 ms, repetition time 56 ms, flip angle 90*^0^*, field-of-view 256 × 256 × 140mm, voxel size 0.5 × 0.5 × 5mm). Freesurfer (Fischl (2012), https://surfer.nmr.mgh.harvard.edu/) was used to preprocess the MRIs, including non-uniform intensity correction and normalisation, and reslicing to an isotropic 1mm volume with field-of-view 256mm in all directions.

Subsequently SPM12 (https://www.fil.ion.ucl.ac.uk/spm/software/spm12/) and Field-trip (Oostenveld et al., 2011) were used to perform non-linear normalisation to the MNI152 template space, and then inverse normalise template brain, skull, and scalp surface meshes, template electrode locations, and template source grids into individual space. The template brain, skull, and scalp meshes were from the ‘standard bem’ head model included in Fieldtrip, which were derived from the Colin27 template MRI (https://www.bic.mni.mcgill.ca/ServicesAtlases/Colin27) as described in Oostenveld et al. (2004). The choice to inverse warp scalp/skull/brain surfaces as opposed to segment each individual MRI and compute individualised surfaces (Ashburner and Friston, 2005) was due to the limited field-of-view of the MRIs, meaning the uppermost points of scalp were often missing (see example in Supplementary Figure S13). Use of template head models for source localisation has been shown to be comparable in quality to individual head models (Fuchs et al., 2002; Henson et al., 2009). The electrode locations were the template ‘standard 1020.elc’ included with Fieldtrip, which are registered to the Colin27 template surface, also inverse warped to individual space.

The template source model consisted of 17 dipoles in regions of interest believed to be associated with SCWT function, namely within the orienting attentional network, executive attention network, and default mode network (Marek et al., 2010; Basten et al., 2011; Rueda et al., 2012; Meier et al., 2012; Duchek et al., 2013; Connolly et al., 2016). Dipole locations were chosen as peak MNI coordinates in previous functional studies of these SCWT-associated resting-state networks (Basten et al., 2011; Schmidt et al., 2013; Conwell et al., 2018). A full table of region names, MNI coordinates, and citations are given in Table S2.

Following nonlinear warping of template brain/skull/scalp surfaces, template electrode locations, and template source models, we computed a 3-layer boundary element method volume conductor model (Fuchs et al., 2002) and the associated leadfield matrix using Fieldtrip. An example warped volume conductor and electrode placement is shown in Supplementary Figure S13. For participants who did not undergo MRI (*n* =1 AD participant, *n* =2 MCI_c_, *n* =4 MCI_nc_, and all HOAs) the same template forward model was used without the nonlinear warping step.

Source localisation was performed using unit-noise-gain LCMV beamforming (Tait et al., 2021; Westner et al., 2022) implemented in Fieldtrip. Connectivity analysis used leakage-corrected amplitude envelope correlations as described in Colclough et al. (2016). In brief, we filtered the data into the frequency band of interest, ran the beamformer source estimation, used symmetric multivariate orthogonalisation to correct for source leakage (Colclough et al., 2015), computed the Hilbert transform to obtain amplitude envelopes, low-pass filtered amplitude envelopes at 1 Hz, resampled the envelopes at 2 Hz, and then computed correlation between each pair of envelopes. This methodology has been shown to give highly reliable connectivity metrics and is robust to noise (Colclough et al., 2016; Tait et al., 2021).

### 2.6 Statistical analysis and GLM

Statistical analyses were performed using MATLAB’s Statistics and Machine Learning Toolbox. Due to the relatively low sample size in this study, in addition to test statistics and p-values, we report Bayes Factors (BF) where possible. To calculate the BF, we used the BayesFactor tool-box (https://klabhub.github.io/bayesFactor/) included with Fieldtrip. Since BFs compare model evidence of null and alternative hypotheses, they have the advantage of differentiating non-significant results due to absence of evidence (e.g. due to small sample size, in which case *BF ∼* 1) and evidence of absence (in which case *BF <<* 1).

To interpret the extent of the relationship between the SCWT-E score and the cluster assignment with the trajectory of MCI subjects (i.e. MCI_c_ or MCI_nc_), we built a statistical model on conversion from MCI to AD. Specifically, we used a generalized linear model (GLM) to model conversion to AD within 2 years. We used GLMs with a logistic link function and binomially distributed response variables, implemented with MATLAB’s `fitglm` function. We fit three GLMs in total, one with the SCWT-E score as a discrete predictor, one with the cluster assignment as a categorical predictor and one with both the SCWT-E score and the cluster assignment as predictors. We subsequently assessed the predictive performance of the GLM by comparing the values obtained from the model to the known outcomes and calculating the area under the receiver operating characteristic (ROC) curve, with confidence intervals (CIs) calculated through bootstrapping sampling.

## 3 Results

### 3.1 Participant demographics

Participant demographics, including sex, age (at initial baseline assessment) and years of education, are outlined in Table 1. Of the 20 MCI participants, 10 converted to AD within two years (two of whom converted within one year) and 10 remained stable after both the one- and two-year follow-ups. No significant differences were observed between healthy older adults (HOAs), people with stable MCI (MCI_nc_), people with MCI converting to AD (MCI_c_), and AD cohorts in terms of age (*F* = 1.93, *p* = 0.141, *BF* = 0.410), education level (*F* = 2.68, *p* = 0.0612, *BF* = 0.883), or sex (*x*^2^ = 2.40, *p* = 0.495). All HOAs underwent the MMSE to rule out cognitive impairment, and all scored 29-30 except one person who scored 27, signifying normal general cognitive health in the HOA cohort.

**Table 1:** Participant demographics.

| Cohort | HOA | AD | MCI<sub>nc</sub> | MCI<sub>c</sub> | ANOVA<br>(F/p/BF) | Chi-squared<br>($\chi^2/p$) | MCI<sub>nc</sub><br>MCI<sub>c</sub><br>vs<br>t-test<br>(t/p/BF) |
| --- | --- | --- | --- | --- | --- | --- | --- |
| N | 11 | 10 | 10 | 10 |  |  |  |
| Male (Fe-male) | 7 (4) | 3 (7) | 5 (5) | 5 (5) |  | 2.40/<br>0.495 |  |
| Age, mean (SD; years) | 70 (5.5) | 72 (4.6) | 73 (3.6) | 75 (5.5) | 1.934/<br>0.141/0.410 |  | 1.11/<br>0.281/0.464 |
| EDU, mean (SD; years) | 8.8 (5.4) | 6.4 (1.7) | 6.1 (1.4) | 5.1 (1.9) | 2.68/<br>0.0612/0.883 |  | 1.34/<br>0.196/0.562 |
| MMSE, mean (SD) | 29.1 (0.83) | N/A | N/A | N/A |  |  |  |

Neuropsychological assessments verified the cognitive status of each group. There was a highly significant effect of diagnosis (HOA vs AD vs MCI) for RAVLT-IR scores (*F* = 37.1, *p* = 1.15 × 10*^-^*^9^, *BF* = 9.23 × 10^6^), RAVLT-DR scores (*F* = 106, *p* = 2.77 × 10*^-^*^16^, *BF* = 2.01 × 10^13^), MFTC-T (*F* = 7.54, *p* = 0.00170, *BF* = 18.8) and MFTC-A (*F* = 7.93, *p* = 0.00130, *BF* = 23.8). As anticipated, trends followed those expected with the HOAs group exhibiting the highest performance, followed by the MCI group, while the AD group demonstrated the lowest performance.

### 3.2 Comparison of neuropsychological testing

As detailed in subsection 2.2, during the baseline assessment, patient cohorts (MCI and AD) underwent an extensive battery of neuropsychological testing to measure cognitive functions associated with disease. These tests were repeated on MCI participants during the follow-up assessments. To better understand the cognitive markers present in patients with MCI that might indicate impending conversion to AD, we retrospectively compared the baseline neuropsychological data between MCI_nc_ and MCI_c_ participants. Figure 2A ranks the BF across all the neuropsychological tests obtained at baseline. BF quantifies the ratio between the likelihood of observing the data under the alternative hypothesis (a difference in means between the cohorts is present) against the likelihood of observing the data under the null hypothesis (no difference in means between the cohorts is present). The SCWT-E was found to have the largest BF, with a value of 4.91. The MCI group that converted to AD (MCI_c_) had higher SCWT-E values at baseline. Supplemental Figure S1 ranks the Cohen’s d effect size across the neuropsychological tests and shows that between MCI_nc_ and MCI_c_ subjects, the largest effect (*d* = 1.14) occurred in the SCWT-E scores. SCWT-E scores can be seen for all subjects in Figure 2B, where, compared to MCI_nc_, the higher SCWT-E scores observed in the MCI_c_ cohort are more similar to the SCWT-E scores found in the AD cohort. We quantified this through a Mann-Whitney U test and BF to compare MCI_nc_ vs AD (*U* = 3.19, *p* = 0.00140, *BF* = 17.6) and MCI_c_ vs AD (*U* = 2.12, *p* = 0.0339, *BF* = 2.84).

**Figure 2:**
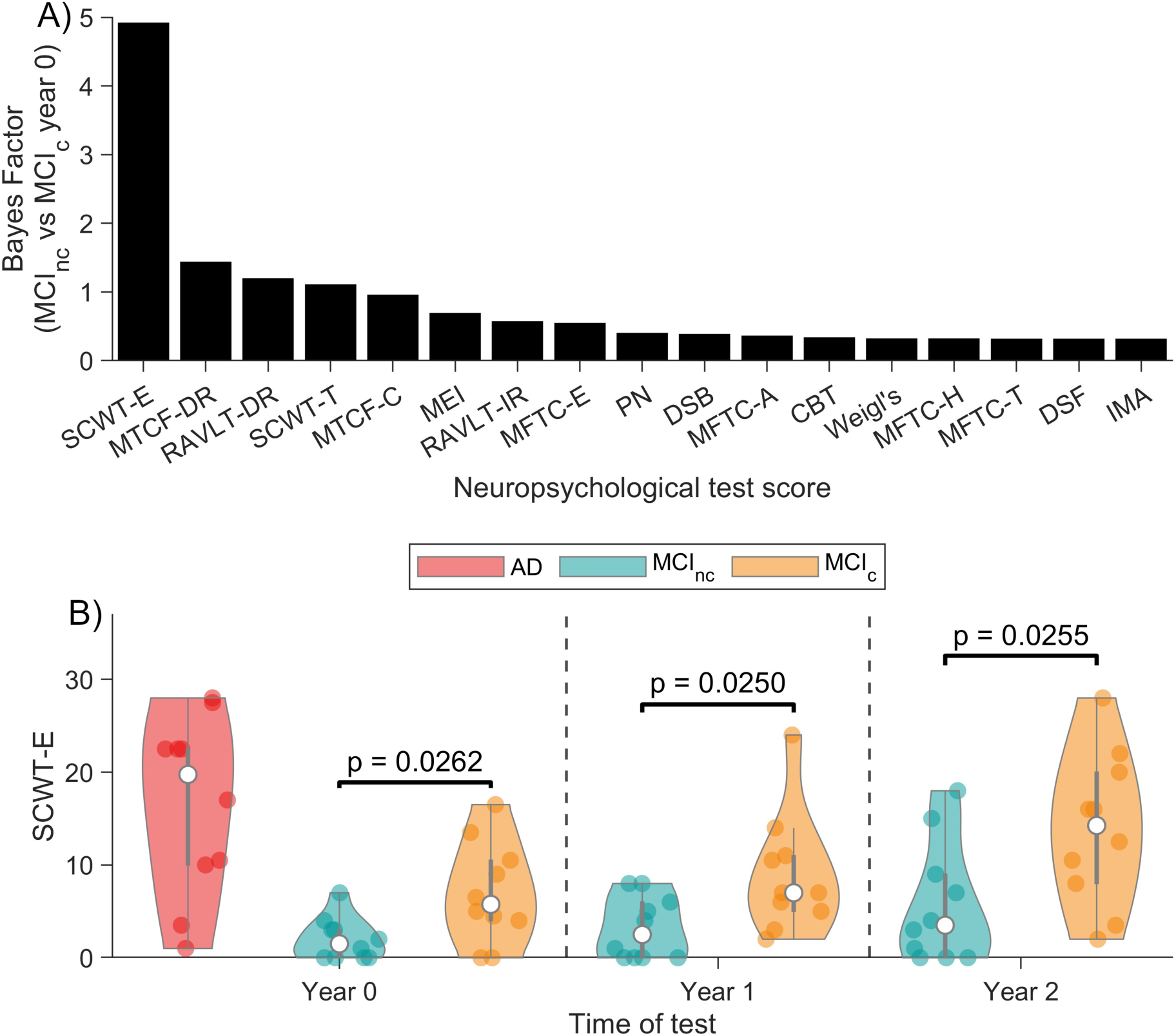
Comparison of neuropsychological test scores between MCI_nc_ and MCI_c_ subjects. A) BF for each neuropsychological test undertaken at baseline between MCI_nc_ and MCI_c_ subjects. The largest BF was observed in the SCWT-E. B) The SCWT-E score between MCI_nc_ and MCI_c_, for years 0, 1 and 2. The SCWT-E for AD subjects is also shown for reference. P-values obtained from a Mann-Whitney U test are shown, indicating that significant differences in SCWT-E scores were observed during each year of assessment. Note, details of the neuropsychological tests used, including the full names, are provided in subsection 2.2.

Figure 2B additionally shows how the SCWT-E scores change across each year of assessment, while Supplemental Figure S2 shows gradients across time. Using a linear mixed effects model for the MCI participants, we found a significant main effect of time (*F* = 11.6, *p* = 6.00 × 10*^-^*^5^) and between-subject effect of conversion status (*F* = 4.13, *p* = 0.0470), but no significant interaction between conversion status and time (*F* = 1.28, *p* = 0.287). Furthermore, BF analysis revealed that the model without interaction term was 4.00 times more likely than the model with interaction. Together, these results suggest that the difference in SCWTE between MCI_c_ and MCI_nc_ does not vary significantly with time, meaning we likely gain as much discriminatory information at time point zero as at follow-ups. This was validated using a non-parametric approach, namely by computing the linear separability between converters and non-converters using the ROC-AUC statistic (a normalised version of the Mann-Whitney U statistic scaled from 0-1) at each time point. At the initial time point, we found AUC=0.795 (95% CI=0.492 - 0.983), while at the initial follow-up the AUC=0.800 (95% CI=0.520 - 0.950) and final time point AUC=0.800 (95% CI=0.541 - 0.950). The overlap in confidence intervals further verifies that there is no significant difference in terms of linear separability of MCI_c_ and MCI_nc_ across time.

### 3.3 Comparison of electrophysiology

The PSD obtained from the EEG data (see subsection 2.4 for details) for each cohort (HOA, MCI and AD) at baseline, and the relative contributions of the delta, theta, alpha and beta frequency bands, were calculated and are given in Supplemental Figure S3. In each case, these PSDs, and the differences observed between cohorts, closely align with the results of previous studies comparing eyes closed resting state EEG between HOAs and patients with AD (Jeong, 2004; Dauwels et al., 2010; Tait et al., 2019), and for patients with MCI (Schumacher et al., 2020; McBride et al., 2014). In particular, spectral slowing and a significant increase in the theta band power were observed in AD compared to HOAs (*U* = 2.23, *p* = 0.0221, *BF* = 5.08), with MCI particpants showing only a subtle non-significant degree of slowing compared to HOAs (Supplemental Figure S3). Furthermore, the PSD and frequency bands obtained by splitting the MCI group by future conversion (i.e. MCI_nc_ and MCI_c_) are given in Supplemental Figure S4. Here, we found no significant differences between the cohorts across any of the frequency bands. However, we did observe that subjects from the MCI cohort (both MCI_nc_ and MCI_c_) appear to have a large degree of variability in their PSD, especially in the alpha frequency band, with a standard deviation across subjects of the relative alpha frequency component to be 0.168. Supplemental Figures 5-8 show the individual PSDs across HOA, AD, MCI_nc_ and MCI_c_ participants, respectively.

To further characterise the electrophysiological data, and to avoid arbitrarily focusing on a particular dominant EEG metric or frequency band, we mapped the full broadband PSD from each participant into 2 dimensions using multi-dimensional scaling (see subsection 2.4 for details). Figure 3A shows a scatter plot of points in this 2-dimensional space. Here, each point represents a participant and distances between points are representative of the differences between the participant’s corresponding PSD (defined by the sum of squares error between spectra across 2-30 Hz). By evaluating the AIC and BIC, we found participants to be congregated into 3 clusters using a Gaussian mixture model (GMM). For further information on cluster assignment, see Supplemental Figure S9. The 3 clusters are shown in Figure 3B, with example spectra from each cluster shown in Figure 3D. Additionally, Figure 3C shows the proportion of participants from each cohort that reside in each cluster. Notably, HOAs were found to reside exclusively in cluster 1. Cluster 1 also contained 90% of the MCI_nc_ participants. Conversely, the majority of AD participants were found to reside in cluster 2, with a high proportion of MCI_c_ participants also in clusters 2 and 3. These differences in cluster membership indicate that EEG PSD analysis can be highly specific to measuring disease progression. The clusters therefore have the potential to help provide a valuable and quantifiable characteristic of pathological changes occurring in the brain during (prodromal) AD.

**Figure 3:**
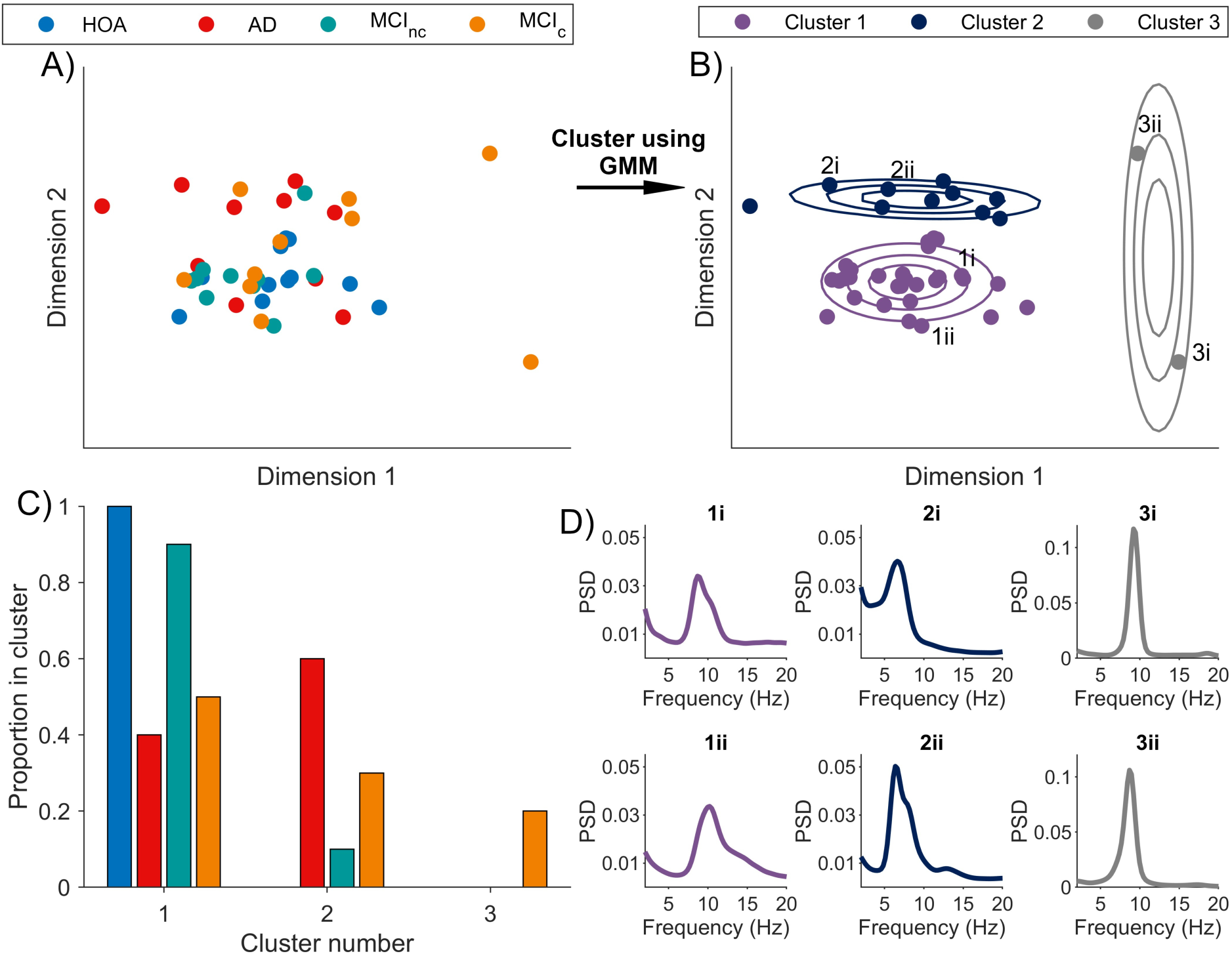
Clustering differences in PSD across subjects. A) Scatter plot of all subjects. Each point represents a subject and distances between points are representative of the differences observed between their corresponding PSD. B) Output from mapping these points into 3 clusters using a GMM. The contour lines denote 25%, 50% and 75% of the multivariate Gaussian distribution for each cluster. C) Proportion of each cohort in each cluster. D) Subplots with two example PSDs from each cluster. Titles of the subplots in D) relate to a subject in the scatter plot labelled in B).

### 3.4 Combining electrophysiological and neuropsychological markers in a GLM

Next, we aimed to combine the observations across neurophysiological testing and electrophysiology to model the progression of prodromal AD. In particular, we built a GLM on the data from MCI_nc_ and MCI_c_ subjects at baseline. Note, in the case of the PSD clustering, membership in cluster 1 is compared against membership in a different cluster. Figure 4 shows that combining the SCWT-E and PSD clustering measures into a concurrent GLM increased the area under the ROC curve to 0.880 (Figure 4C), compared to either 0.795 (SCWT-E; Figure 4B) or 0.710 (PSD clustering; Figure 4A) when a single variable was used. A permutation test verified these effects, with the GLM from the SCWT-E, PSD clustering and SCWT-E/PSD clustering combined obtaining a p-value of 0.0260, 0.130 and 0.0180, respectively. Hence, the integration of multiple functional markers into a statistical test can be useful in characterising the progression of prodromal AD.

**Figure 4:**
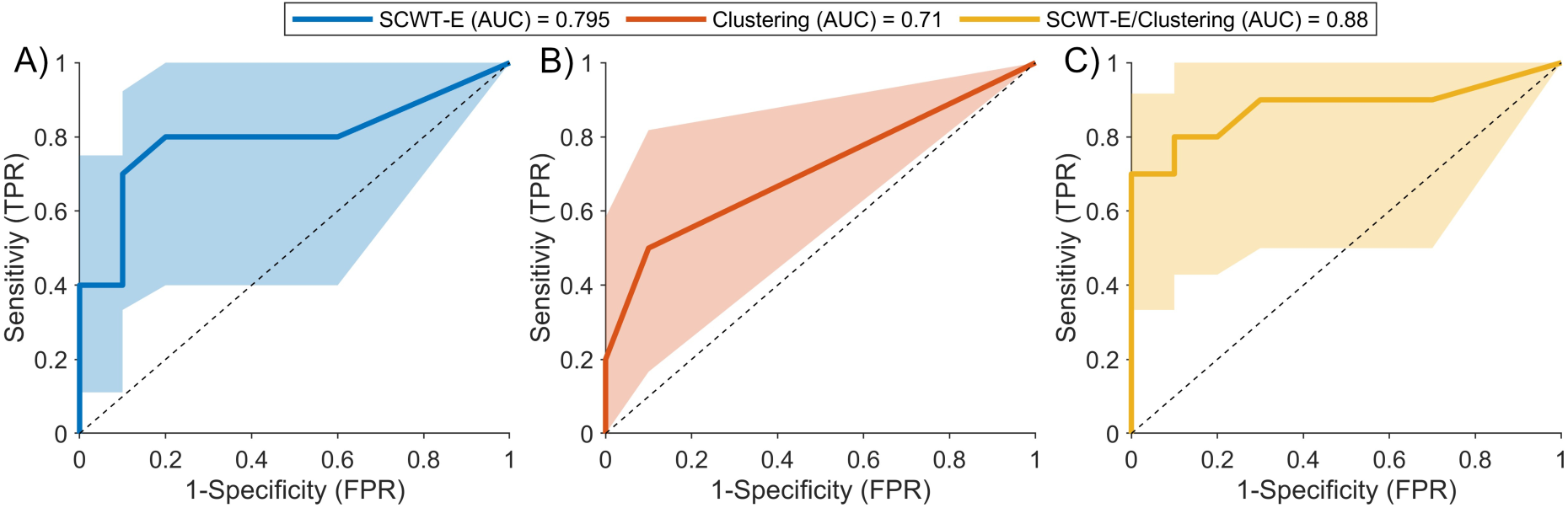
ROC curves for differentiating between MCI_nc_ and MCI_c_ subjects. A) ROC curve using the SCWT-E score as a predictor, obtaining an AUC of 0.795 (95% CI = [0.495,0.960]). B) ROC curve using the EEG clustering score as a predictor, obtaining an AUC of 0.710 (95% CI = [0.510,0.875]). C) ROC curve using the combined SCWT-E and clustering as predictors, obtaining an AUC of 0.880 (95% CI = [0.560,0.0.995]). In each case, the mean and 95% CIs are shown by the solid lines and shaded regions, respectively.

### 3.5 Interpreting SCWT-E deficits in patients with prodromal AD

Thus far, we have characterised neurophysiological tests and electrophysiological properties. In both these domains, we observed differences between MCI_nc_ and MCI_c_ subjects at baseline. We then found that a GLM comprised of combining these measures had the greatest AUC, indicating a degree of orthogonality between them. We next aimed to understand the mechanisms driving these differences. First, we explore the network mechanisms that underline the SCWT-E scores.

Performance in the SCWT is believed to be associated with connectivity in cortical resting-state networks associated with working memory and attention, including the orienting attentional network, executive attentional network, and default mode network (Marek et al., 2010; Basten et al., 2011; Rueda et al., 2012; Meier et al., 2012; Duchek et al., 2013; Connolly et al., 2016). Hence, we hypothesised that impaired SCWT-E scores in people with AD (at either dementia or prodromal - i.e. MCI_c_-stage) are associated with disconnectivity in these resting-state networks. To test this, connectivity within the orientating attentional, executive attentional, and default-mode networks was calculated by combining electrophysiological dynamics from EEG with anatomical information from MRI using a beamforming approach (subsection 2.5). Mean connectivity strength across all three subnetworks was significantly negatively correlated with SCWT-E score in the theta (4-8 Hz, *R* = −0.597, *p* = 0.00630, *BF* = 7.79, Figure 5A) and beta (13-30 Hz, *R* = −0.444, *p* = 0.0497, *BF* = 1.16, Figure 5D) bands, while similar but non-significant negative correlations were observed in alpha (8-12 Hz) and gamma (30-70 Hz) bands (Figure S12). These correlations were calculated using robust linear regression. These results suggest broadband disconnectivity in the working-memory and attention networks are associated with more errors in the SCWT task, with the strongest associations occurring across theta and beta frequencies. To gain more anatomical information, we tested connectivity within each of the three subnetworks. In the theta band, the negative correlation between mean connectivity strength and SCWT-E score remained significant for the orienting attention (*R* = −0.516, *p* = 0.0230, *BF* = 2.53) and executive control (*R* = −0.491, *p* = 0.0288, *BF* = 1.88) subnetworks, and exhibited a negative but non-significant correlation in the default-mode network (*R* = −0.431, *p* = 0.0667, *BF* = 1.02). Conversely, in the beta band, the default-mode network exhibited the strongest effect (*R* = −0.505, *p* = 0.0233, *BF* = 2.20), while the two attention networks were non-significant (orienting: *R* = −0.370, *p* = 0.109, *BF* = 0.615; executive: *R* = −0.385, *p* = 0.935, *BF* = 0.691). Thus, our results suggest that impaired connectivity at slower frequencies within the orienting and executive attention networks, combined with impaired connectivity at faster frequencies within the default-mode network, are potentially associated with reduced SCWT-E scores in people with (prodromal) AD.

**Figure 5:**
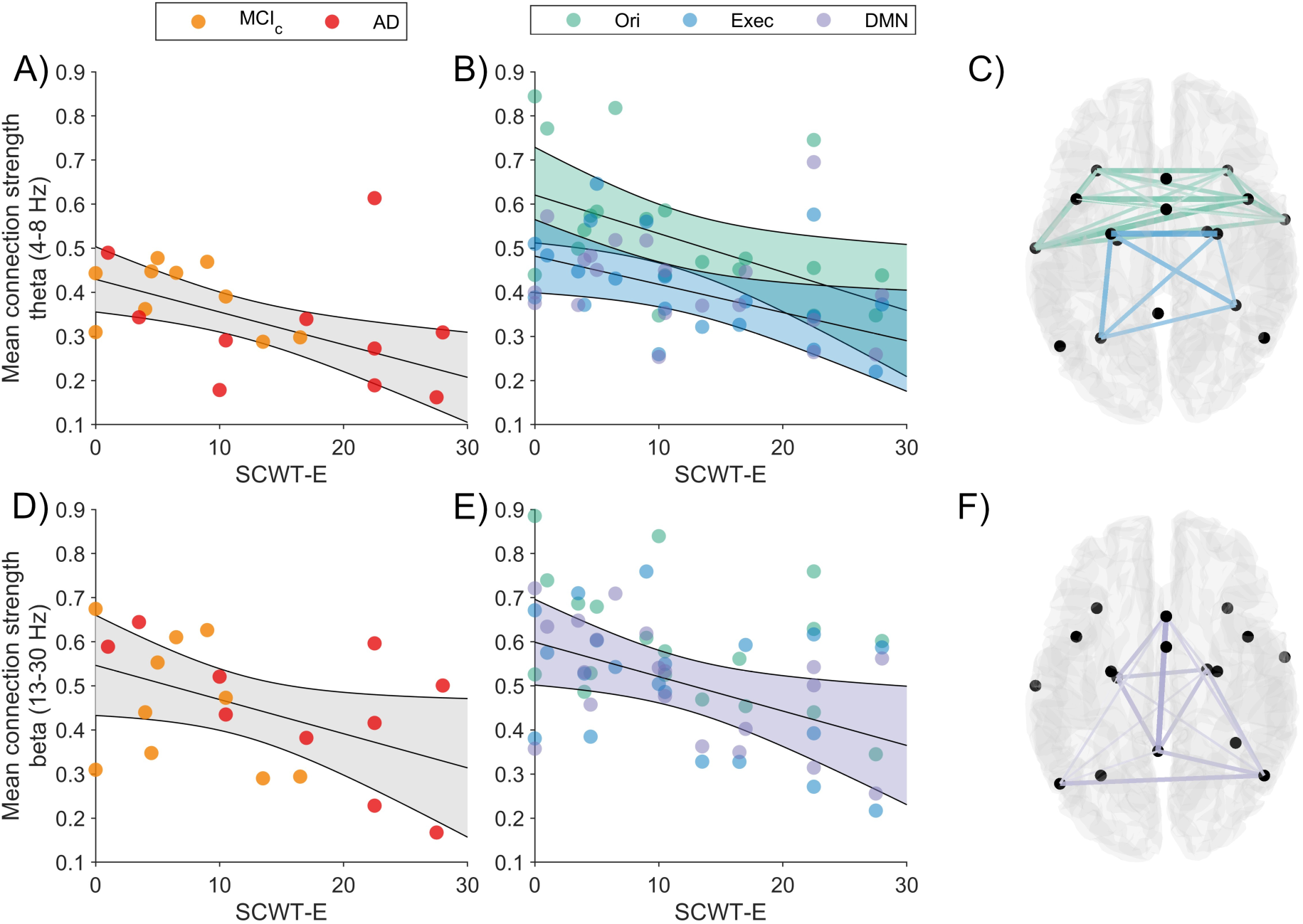
Degree of impairment to SCWT-E scores in people with (prodromal) AD is correlated with functional disconnectivity. In people with (prodromal) AD (i.e. the AD or MCI_C_ groups), mean connection strength across the 17 nodes of the SCWT attention/working-memory network (see subsection 2.5) significantly negatively correlates with SWCT-E in the A) theta and D) beta bands. This correlation is significant within the orienting (Ori) and executive (Exec) attention subnetworks for the B) theta band, and default-mode network (DMN) for the E) beta band. In C), we show the spatial distribution of the 17 nodes, and connections making up the Ori (green) and Exec (orange) networks. Edge weights are proportional to the strength of correlation between the theta-band amplitude envelope correlation of that edge and SCWT-E score. F) gives the same as in C), but for beta-band DMN edges.

### 3.6 Interpreting PSD clusters using mathematical modelling

Finally, we used mathematical modelling to interpret the PSD clustering results. First, we characterised differences in the spectral clusters by applying spectral parameterisation (Donoghue et al., 2020). Differences between clusters were primarily in the oscillatory component as opposed to the aperiodic component, and were associated with theta-alpha band oscillations (Supplemental Figure S10). Furthermore, we found that cluster 1 mainly contained typical eyes-closed resting alpha activity peaking at approximately 9-10 Hz, along with subjects that had no dominant alpha rhythm peak (i.e. 1/f spectrum). Cluster 2 was characterised by a slowing of the alpha rhythm to approximately 8 Hz, which is reminiscent of a pathological state (Smith, 2005), and cluster 3 contained significantly increased peak alpha power. The spectral parameterisation also revealed that there were significant differences in the peak frequency across clusters (Supplemental Figure S10F) and that the aperiodic slope is indifferent across clusters (Supplemental Figure S10E), in line with previous studies suggesting the aperiodic slope is not important in neurodegenerative disease (Kopcanova et al., 2024).

While spectral parameterisation is useful as descriptive modelling, i.e. to characterise and quantify differences in the spectrum, and to gain some mechanistic insight (periodic oscillations vs aperiodic noise), it does not quantify spectral profiles in terms of synaptic or cellular mechanisms. We therefore subsequently used a generative mathematical model to infer the synaptic mechanisms underlying the difference observed. In particular, we optimised the parameters from the Liley neural mass model (D T J Liley and Dafilis, 2002) to properties of the EEG using a previously developed multi-objective evolutionary algorithm (MOEA; Dunstan et al. (2023)). For details on the neural mass model and optimisation algorithm used, including a description of parameter values and model equations, see Supplemental 4. In short, neural mass models are well suited to model the activity recorded on EEG and have extensively been used to interpret the synaptic mechanisms that generate various healthy and pathological brain rhythms (David and Friston, 2003), including those observed in AD (for example, see Bhattacharya et al. (2011)). Neural mass models describe the synaptic interactions between populations of excitatory and inhibitory neurons. The mechanistic understanding we can obtain from them includes the time scales and amplitudes of post-synaptic membrane potentials at the level of activity in each population (their firing rates).

Figure 6A shows the PSD data (mean and standard error across subjects) for each of the clusters. In contrast, Figure 6B shows the PSD (mean and standard error across subjects) from model simulations after using the MOEA to fit the model to the baseline EEG data. It can be seen that the simulations accurately recreate the PSD and differences observed across the clusters. Supplemental Figures S14, S15 and S16 show the distribution of parameters recovered from using the MOEA to fit the data from subjects in clusters 1, 2 and 3, respectively. Figure 6C and D show the mean gain (amplitude) of the postsynaptic potentials (PSPs) in the neural mass for the excitatory and inhibitory populations (i.e. excitatory PSP and inhibitory PSP), respectively. Furthermore, Figure 6E and F show the mean neuronal population firing rate curves in the neural mass for the excitatory and inhibitory populations, respectively. The differences in both the firing rates and PSP waveforms obtained from the model indicate that the PSD clusters capture different synaptic physiologies. In particular, we found that the cluster containing the healthy dynamics (cluster 1) is underpinned by higher excitatory PSP amplitudes (with a comparison of mean PSP peak across clusters obtaining *F* = 9.55, *p* = 0.000437, *BF* = 138), faster inhibitory PSP timescales (with a comparison of mean inhibitory timescale parameter γ*_i_* obtaining *F* = 4.06, *p* = 0.0252, *BF* = 3.16) and increased but insignificant propensity for inhibitory neurons to fire (with a comparison of half activation of sigmoid across clusters obtaining *F* = 2.66, *p* = 0.0827, *BF* = 0.812). Thus, the clusters containing the pathological dynamics (clusters 2 and 3) are underpinned by a reduction in excitatory synaptic efficacy and an increase in inhibitory synaptic efficacy, although we found these to be somewhat balanced by a decrease in the activity of inhibitory populations to fire.

**Figure 6:**
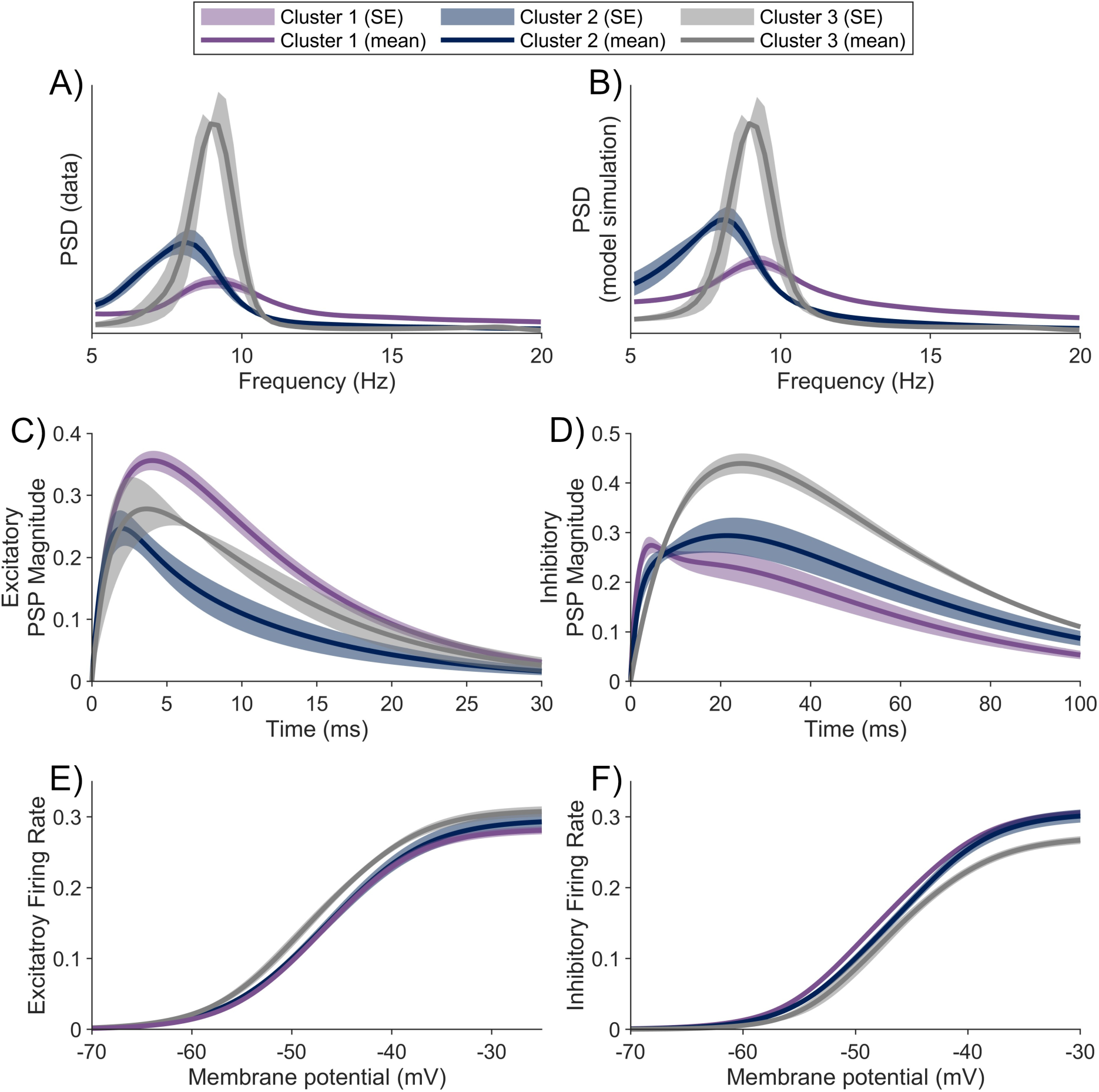
Comparison of neural mass model properties from simulations across PSD clusters. A) and B) show the PSD across clusters from data and model simulations after fitting to data, respectively. The mean PSPs from the model simulations for the C) excitatory and D) inhibitory populations are shown, as are the mean firing rate curves from the E) excitatory and F) inhibitory populations of the model. In each case, the line denotes the mean and the shaded region denotes the standard error (SE) across subjects in the cluster (see legend).

## 4 Discussion

In this study, we analysed EEG data and an extensive battery of neuropsychological tests recorded from people with AD, people with amnestic MCI and HOAs. We followed and tracked the cognitive progression of the MCI group for a further 2 years and retrospectively classified them into converter (MCI_c_) or non-converter (MCI_nc_) subgroups based on whether the criteria for an AD diagnosis had been met within the two-year follow-up period. We observed substantial differences between MCI_c_ and MCI_nc_ at baseline in both behavioural and electrophysiological measures of brain function, namely the number of errors made during the SCWT and the EEG power spectrum. We subsequently explored the network and synaptic mechanisms underlying these differences. We found the SCWT-E scores correlated with functional disconnectivity in the theta and beta frequency bands and, using mathematical modelling, we mapped altered PSDs to underlying synaptic mechanisms. The multi-modal, longitudinal, and interdisciplinary nature of this work presents important progress in understanding AD progression not only by characterising early-stage (prodromal) functional fingerprints of AD - both behavioural and electrophysiological - but also through novel insight into the mechanisms that drive these functional changes.

### 4.1 SCWT-E deficits in prodromal AD

Neuropsychological testing has been established as a useful tool to detect subtle cognitive impairment in patients with AD, often before the onset of common symptoms associated with the disease (Jacobs et al., 1995; Albert et al., 2001; Tabert et al., 2006). In this study, we performed a longitudinal analysis across a wide range of neurophysiological tests to quantify cognitive ability across domains including short-term memory, long-term memory and attentive-executive functions. The SCWT-E scores differentiated the MCI_c_ and MCI_nc_ cohorts by a much greater margin than any other cognitive test performed (BF = 4.91 compared to the next most differential test which had BF = 1.43). Interestingly, our results show that these differences were consistent over time using both parametric (linear mixed effects modelling, BF analysis) and non-parametric (linear separability via ROC-AUC scores) approaches. These results suggest that separating converters from non-converters at the initial MCI stage is indistinguishable from the same analysis at a two-year follow-up, meaning the SCWT-E score has great potential as a possible early-stage diagnostic marker.

The SCWT-E score is associated with executive functioning and attention, processing speed, working memory and general cognitive efficiency and flexibility (see Scarpina and Tagini (2017) for a review). Hence the differences in SCWT-E between MCI_c_ and MCI_nc_ in our cohort likely reflects the widely reported impairments to executive functioning, attention, and working memory in early-stages of Alzheimer’s dementia (Perry and Hodges, 1999; Albert et al., 2001; Baudic et al., 2006; Arvanitakis et al., 2019; Guarino et al., 2019). Indeed, a systematic review of the literature suggests that the SCWT outperforms other measures of executive performance for differentiating AD from healthy ageing, potentially due to the effects of working memory performance on the task (Guarino et al., 2019). The results presented here support that this occurs even at the prodromal stages of the disease, suggesting that the SCWT may be potentially useful as a prodromal marker for AD. Future work should aim to test this in larger cohorts of participants.

The Stroop task is believed to activate cortical networks such as the orienting and executive attention networks and the default mode network, which anatomically consists of nodes in frontal, parietal, and medial temporal areas, as well as cingulate and insular cortices (Fox et al., 2005; Marek et al., 2010; Basten et al., 2011; Rueda et al., 2012; Meier et al., 2012; Duchek et al., 2013; Connolly et al., 2016). These areas also strongly overlap with those forming the cortical working memory network (see Fox et al. (2005) and the BrainMap database’s meta-analysis of 407 publications https://brainmap.org/taxonomy/behaviors/Cognition.Memory.Working.html). Disconnection in the fronto-temporal-parietal networks has widely been reported in early AD and associated with attentional and executive deficits (Neufang et al., 2011; Zhao et al., 2018; Tait et al., 2019; Buzi et al., 2023). Here we demonstrated a significant correlation between SCWT-E scores and disconnection of the orienting attention, executive attention, and default mode networks, supporting the hypothesis that long-range disconnection may drive cognitive impairment in early AD. We further showed that this disconnectivity is specific to early AD and not a marker of general cognitive impairment, as connectivity was reduced only in MCI_c_ and AD relative to controls, and not MCI_nc_ (Figure S11). Interestingly, mounting recent evidence from the EEG microstate literature (Tarailis et al., 2024) suggests these impairments may be related to impaired transitioning of dynamic networks associated with cognition and fronto-parietal/temporal connectivty on a millisecond timescale (Nishida et al., 2013; Smailovic et al., 2019; Schumacher et al., 2020; Musaeus et al., 2019, 2020; Tait et al., 2020; Lian et al., 2021; Tarailis et al., 2024). In this study, only static connectivity was analysed, but further insight may be gained through a dynamic connectivity/microstate analysis. In particular, the electrophysiological mechanisms underpinning microstates and dynamic connectivity are not well understood, nor are the processes in which cognition arises from these dynamic networks, so future work should focus on these mechanisms to better understand cognitive deficits in early AD.

### 4.2 Clusters of electrophysiological spectra characterise prodromal AD

Different patterns of electrophysiological activity in people with AD, compared to HOAs, have been well documented (see Cassani et al. (2018) for a review). One of the most robustly reported differences is a slowing of the EEG frequency spectrum observed in AD (Smith, 2005; Dauwels et al., 2010; Moretti et al., 2011; Benz et al., 2014; Tait et al., 2019; Kopcanova et al., 2024). These spectral changes have also been reported at the prodromal stage as a potential early signature of AD (Poil et al., 2013; Rossini et al., 2006; Moretti et al., 2011). Using traditional frequency-band analyses, we found AD-associated spectral fingerprints which strongly align with those reported in the existing literature, namely a reduced peak frequency of alpha oscillations and increased ratio of slow power in people with AD (Supplemental Figure S3).

Here we took an alternative approach by using unsupervised learning to comprehensively characterise common spectral profiles within our cohort. This approach has several advantages over the traditional frequency-band based analysis. First, there is no need to *a priori* select frequency bands of interest. While EEG frequency bands are well characterised in normative data (Buzsáki et al., 2012), there is variability in these bands across aging (Donoghue et al., 2020; Quinn et al., 2024) and clinical groups (Tait et al., 2019; Kopcanova et al., 2024). Additionally, because this approach is unsupervised, it will not suffer from overfitting and enables the quantification of the disease likelihood in terms of a deviation from a computer-learned normative cluster. This is different from a direct comparison (such as a t-test) which requires a predefined band to separate cohorts and a specific homogeneous change in one direction. The comparison of cluster assignment therefore allowed us to analyse differences in the EEG PSD using an entirely data-driven approach. This analysis revealed that cluster assignment was significantly associated with diagnosis, indicating this method shows potential for an accurate and unbiased approach to deciphering electrophysiological activity across the cohorts, including between MCI_c_ and MCI_nc_.

To interpret the electrophysiological differences captured across the EEG PSD clusters, we additionally used mathematical modelling to delineate neural population activity from the data. Mathematical modelling of large-scale brain activity has become a pivotal tool in neuroscience to elucidate the underlying mechanisms driving the different dynamic patterns observed in disease (Breakspear, 2017). In particular, neural mass models are well suited to investigate the mechanisms at the level of excitatory and inhibitory neural populations that contribute to the understanding of electrophysiological signals, including pathological patterns that manifest in patients with AD. A recent study (Ranasinghe et al., 2022) that optimised parameters of a neural mass model to magnetoencephalographic data showed evidence of abnormal synaptic activity in patients with AD. In particular, their model indicated differences in excitatory and inhibitory time constant parameters between patients with AD and HOAs, and found that these parameters correlated with AD patients’ levels of tau and Ab proteins. In this study, we optimised the parameters of the Liley neural mass model (D T J Liley and Dafilis, 2002) to the baseline EEG data from all the subjects. We then compared emergent properties of the model that were required to recapitulate the EEG data. We note that the Liley model is widely used to study alpha rhythm EEG data and therefore, combined with the optimisation algorithms we used, enabled us to better recreate the PSD from the data compared to previous studies with other neural mass models (such as Ranasinghe et al. (2022)). We found the spectral clusters differed predominantly by the mean synaptic gain and mean synaptic time constants in the model. In particular, clusters that were largely populated by AD and MCI_c_ subjects were found to have a less efficient and faster excitatory postsynaptic potential, but a slower inhibitory postsynaptic potential, indicating that this pipeline is capable of mapping different electrophysiological properties observed in disease to synaptic mechanisms that may be causative for the disease prognosis. This approach has the potential for substantial implications in understanding the manifestation of the prodromal stage of AD and consequently in the future could be used to inform potential treatment protocols.

### 4.3 Limitations and future perspectives

A key limitation of this study is the relatively small sample size of 20 MCI subjects. To help mitigate uncertainty in results due to the limited sample size, we reported BF (as well as p-values) wherever possible, which can help differentiate between evidence of absence and absence of evidence. Thus, with the use of BF one can estimate whether a certain statistic is unchanging between two cohorts, or whether sample size is too small to give useful insight.

Furthermore, due to the limited sample size and absence of a hold-out validation set, here we did not aim to build a predictive classifier for prodromal AD. Crucially, we did not attempt to mine the data for an array of features that can best generate a classifier for predicting the conversion of MCI to AD. Although this approach has generated promise for EEG as an early diagnostic tool for AD (Poil et al. (2013), Jelic et al. (2000), Vecchio et al. (2018)), these methods often involve selecting an arbitrarily high number of features from the EEG data (for example, 177 EEG biomarkers were used in Poil et al. (2013)). Rather, in this study, we aimed to perform a statistical analysis of a small but highly multi-modal longitudinal dataset to uncover potential markers for prodromal AD and understand their mechanistic underpinnings. For example, while we combined neuropsychological and electrophysiological test scores in a logistic regression model to find AUC values of 0.88, one should not interpret this model as a predictive classifier. It is unclear how representative this dataset is of wider cohorts, since features were selected based on the data with no hold-out set on which to test generalisability. Our models were instead used in a non-parametric permutation-based statistical testing framework to assess significance against permutation surrogates. In this regard, AUC should be treated as a test statistic, in line with the univariate case in which AUC is a normalised version of the Mann-Whitney test statistic. Future work should involve developing and testing predictive models on larger cohorts, building on the statistical markers uncovered herein.

The small sample size reported here was a trade-off for a key strength of the study, namely highly descriptive longitudinal data. By following a small group longitudinally, we were able to collect a highly detailed neuropsychological battery, and functional (EEG) and structural (MRI) data. This small, but detailed, dataset should therefore act as a guide for future large-cohort studies, and future work should focus on resting EEG and Stroop testing in a wider group of individuals to build potentially clinically useful predictive models and better understand the underpinning mechanisms of AD.

### 4.4 Conclusions

Our study supports the potential of multi-modal tools, that are inexpensive, non-invasive, and interpretable, for characterising prodromal AD. Understanding the mechanisms that contribute to differences observed could indicate that the measures discussed will be more robust at generalising to other datasets. In this regard, in the future, EEG PSD clustering and the SCWT-E score could be used to test statistical models to predict the accuracy of diagnosis in larger independent studies. This work will therefore ultimately lead to the elucidation of the full potential of combining functional measurements to improve the diagnosis and prognosis of AD.

## Contributions

**DMD**: Conceptualization, Methodology, Formal analysis, Writing - Original Draft, Writing - Review & Editing. **EB**: Conceptualization, Investigation, Writing - Review & Editing. **SG**: Conceptualization, Resources, Writing - Review & Editing, Project administration, Funding acquisition. **RF**: Investigation, Resources. **BV**: Investigation, Resources. **MV**: Investigation, Resources. **MC**: Investigation, Resources. **CM**: Conceptualization. **FT**: Conceptualization, Writing - Original Draft, Writing - Review & Editing, Supervision, Project administration, Funding acquisition. **MG**: Conceptualization, Methodology, Writing - Original Draft, Writing - Review & Editing, Supervision, Funding acquisition. **LT**: Conceptualization, Methodology, Formal analysis, Writing - Original Draft, Writing - Review & Editing, Supervision

## Notes

### Competing Interest Statement

The authors have declared no competing interest.

