## Supplementary Material for "Longitudinal assessment of the conversion of mild cognitive impairment into Alzheimer’s dementia: Observations and mechanisms from neuropsychological testing and electrophysiology"

### Supplemental

#### Supplemental 1: Neuropsychological testing

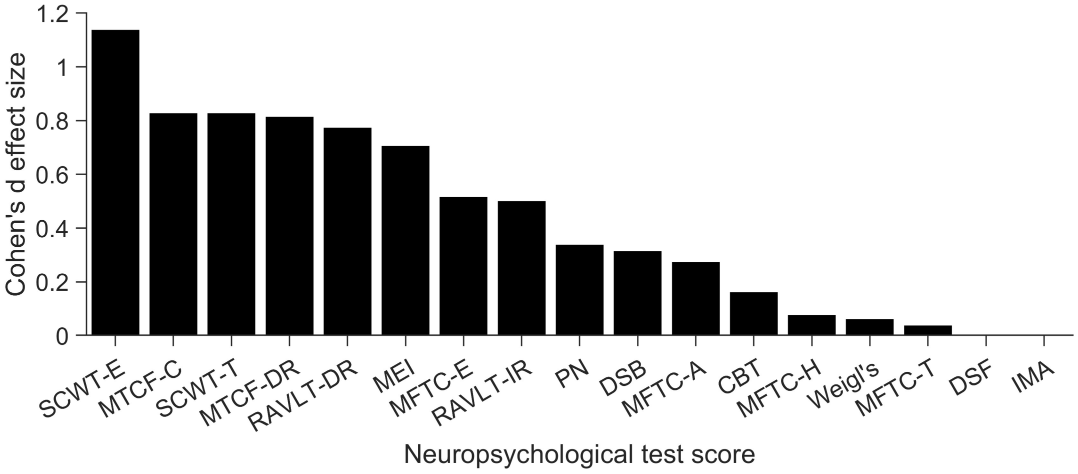


Supplemental Figure S1: **Cohen's d effect size comparison of neuropsychological test scores between $MCI_{nc}$ and $MCI_c$ subjects.**

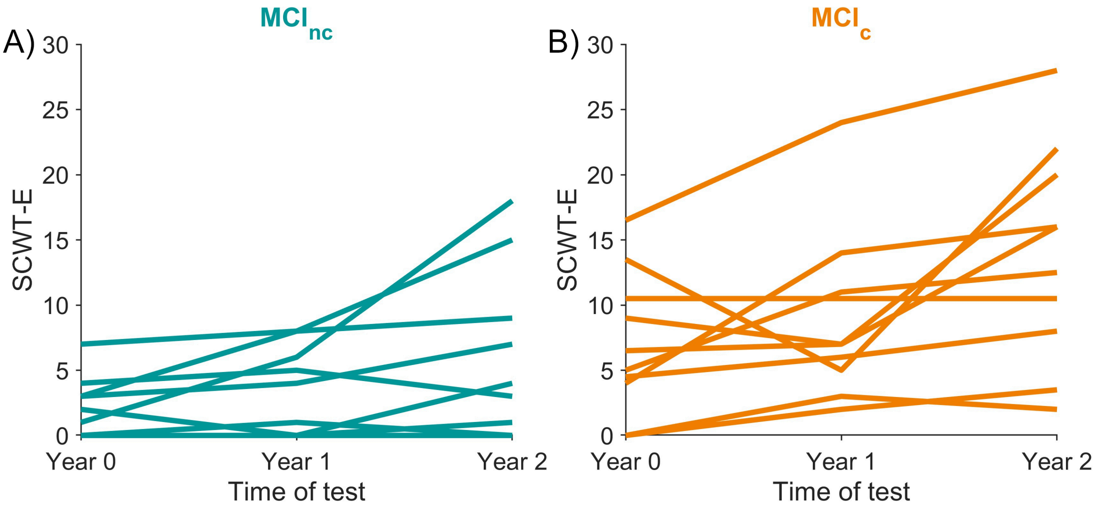


Supplemental Figure S2: **SCWT-E score comparison for MCI subjects overtime.** Both the A) $MCI_{nc}$ and B) $MCI_c$ subjects in general show an increasing trend of SCWT-E scores overtime.

#### Supplemental 2: Power spectral density

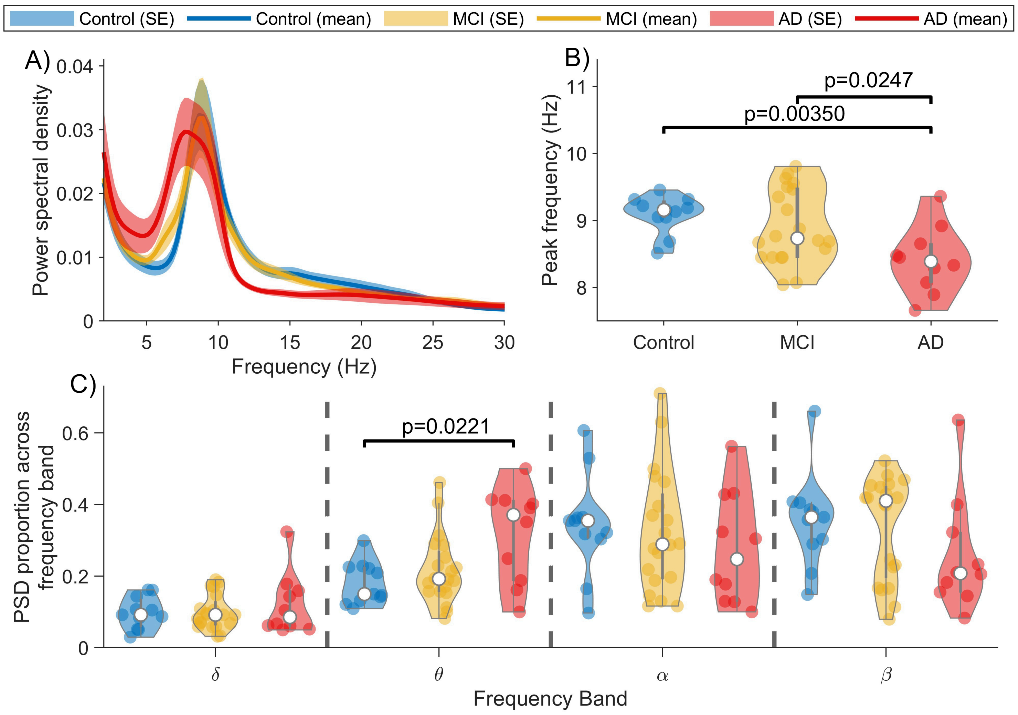


Supplemental Figure S3: **Spectral slowing is present in patients with AD.** A) PSD waveforms for control, MCI and AD cohorts. The line denotes the mean across subjects, and the shaded area represents standard error from within each cohort. B) Peak frequency across the theta-alpha band. AD patients showed a significant reduction in peak frequency compared to controls ($U = 2.92$, $p = 0.00350$, $BF = 16.0$) and MCI ($U = 2.25$, $p = 0.0247$, $BF = 0.809$). C) Relative PSD in each frequency band for each cohort. AD patients showed a significant increase in $\theta$ power compared to controls ($U = 2.29$, $p = 0.0221$, $BF = 5.08$)

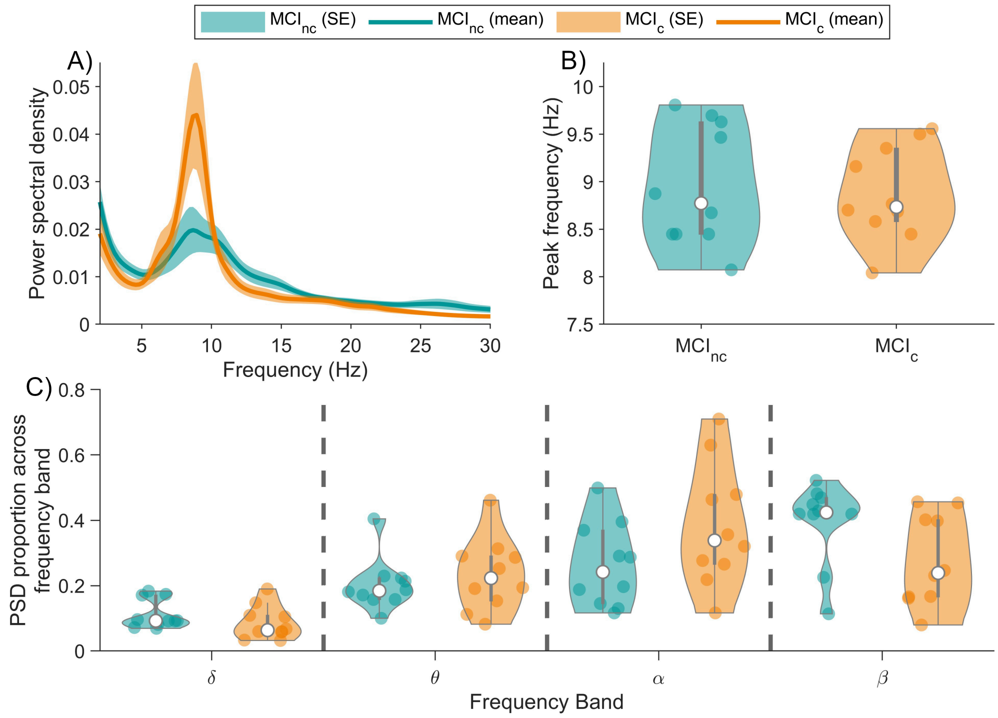


Supplemental Figure S4: **PSD waveforms between $MCI_c$ and $MCI_{nc}$.** A) PSD waveforms for the $MCI_c$ and $MCI_{nc}$ cohorts. The line denotes the mean across subjects, and the shaded area represents standard error from within each cohort. B) Peak frequency across the theta-alpha band. C) Relative PSD in each frequency band for each cohort. No significant differences were observed in peak frequency or a particular frequency band.

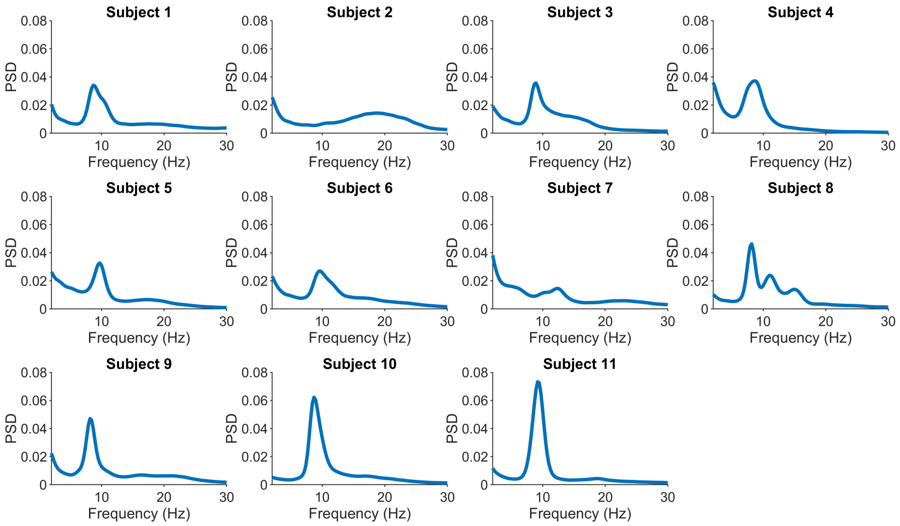


Supplemental Figure S5: **PSD from each control subject.**

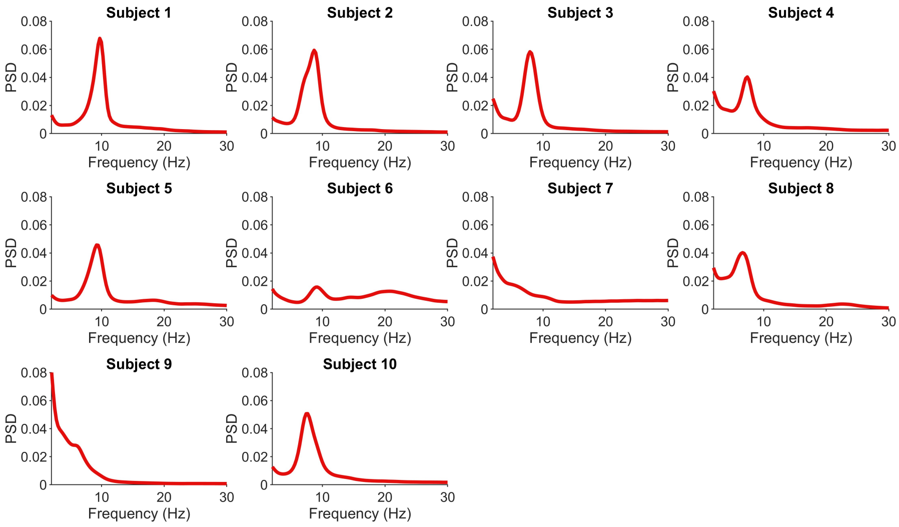


Supplemental Figure S6: **PSD from each AD subject.**

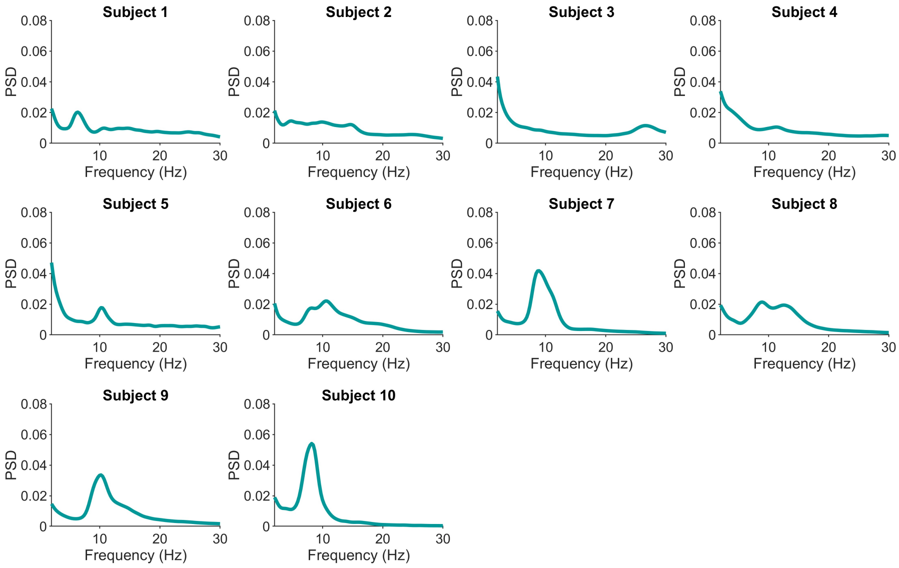


Supplemental Figure S7: **PSD from each MCI$_{nc}$ subject.**

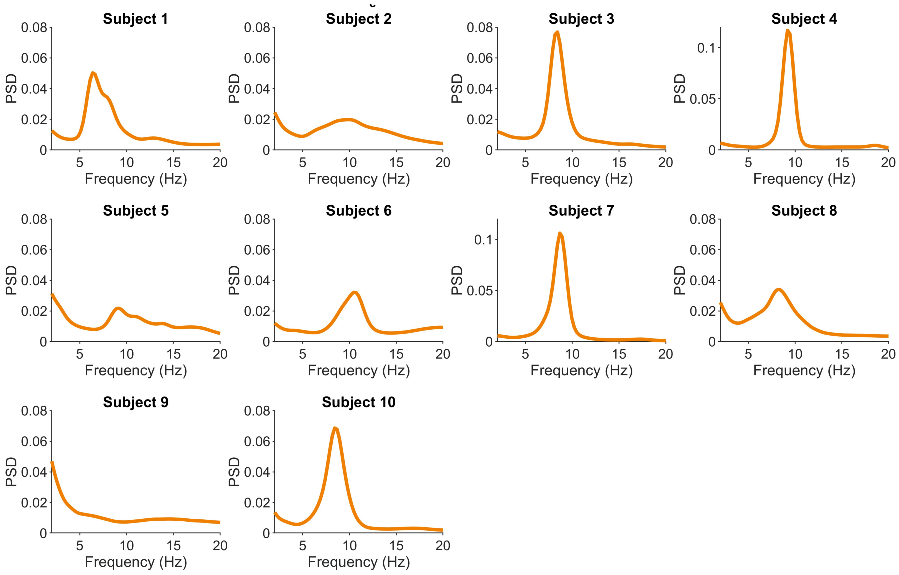


Supplemental Figure S8: **PSD from each MCI$_c$ subject.**

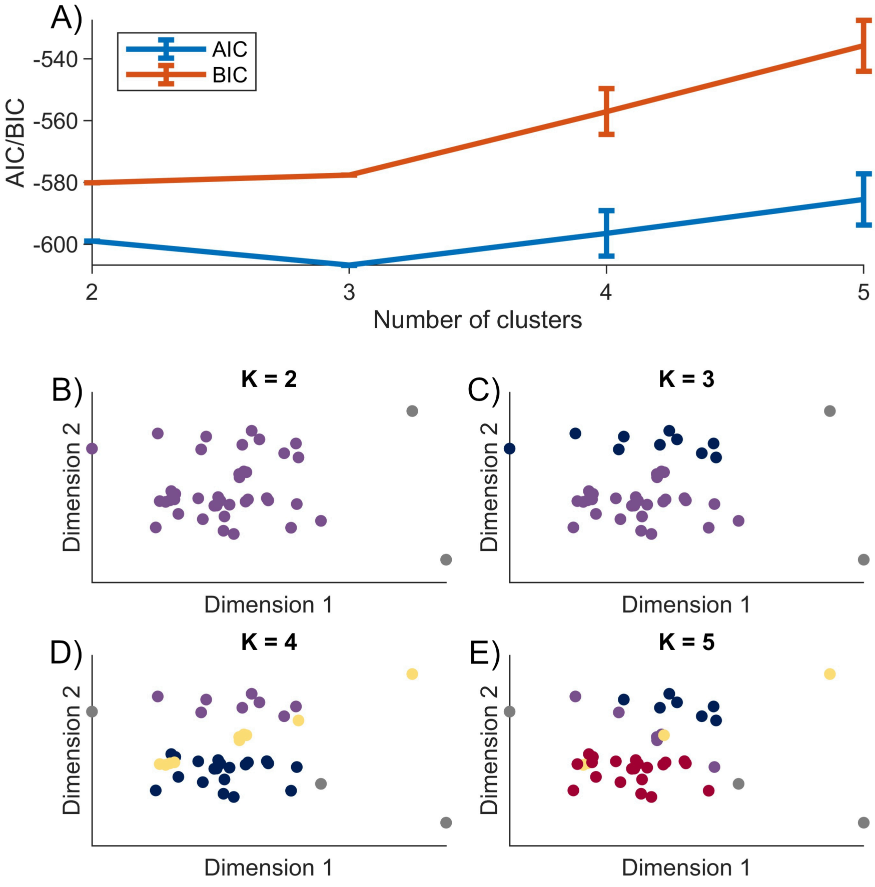


Supplemental Figure S9: **K=3 clusters were selected using a Gaussian mixture model.** A) AIC and BIC performance for k=2,3,4 and 5 clusters using a Gaussian mixture model on the embedded PSD distance matrix. B-D) shows an example of the cluster assigned (each colour represents the cluster membership of each point) for k=2, 3, 4 and 5, respectively.

To characterise differences in spectra across clusters, we used spectral parameterisation (also called fitting oscillations and one-over-f or FOOOF; Donoghue et al. (2020)) applied to each individual's average (across channels) PSD. We used Brainstorm's implementation of spectral parameterization (Tadel et al., 2011), using the default parameters. Spectral parameterisation separates the PSD into aperiodic one-over-f type components and periodic oscillatory bumps in the spectrum, essentially allowing the user to characterise an aperiodic spectrum and a periodic spectrum. Previous work has suggested that spectral slowing in AD is associated primarily with the oscillatory spectrum (Azami et al., 2023; Kopcanova et al., 2024). In contrast, the slope of the aperiodic component has been associated with excitatory-inhibitory balance (Gao et al., 2017). Therefore, spectral parameterisation may give insight into the mechanisms underpinning differences observed in the PSD between clusters, and their relation to disease.

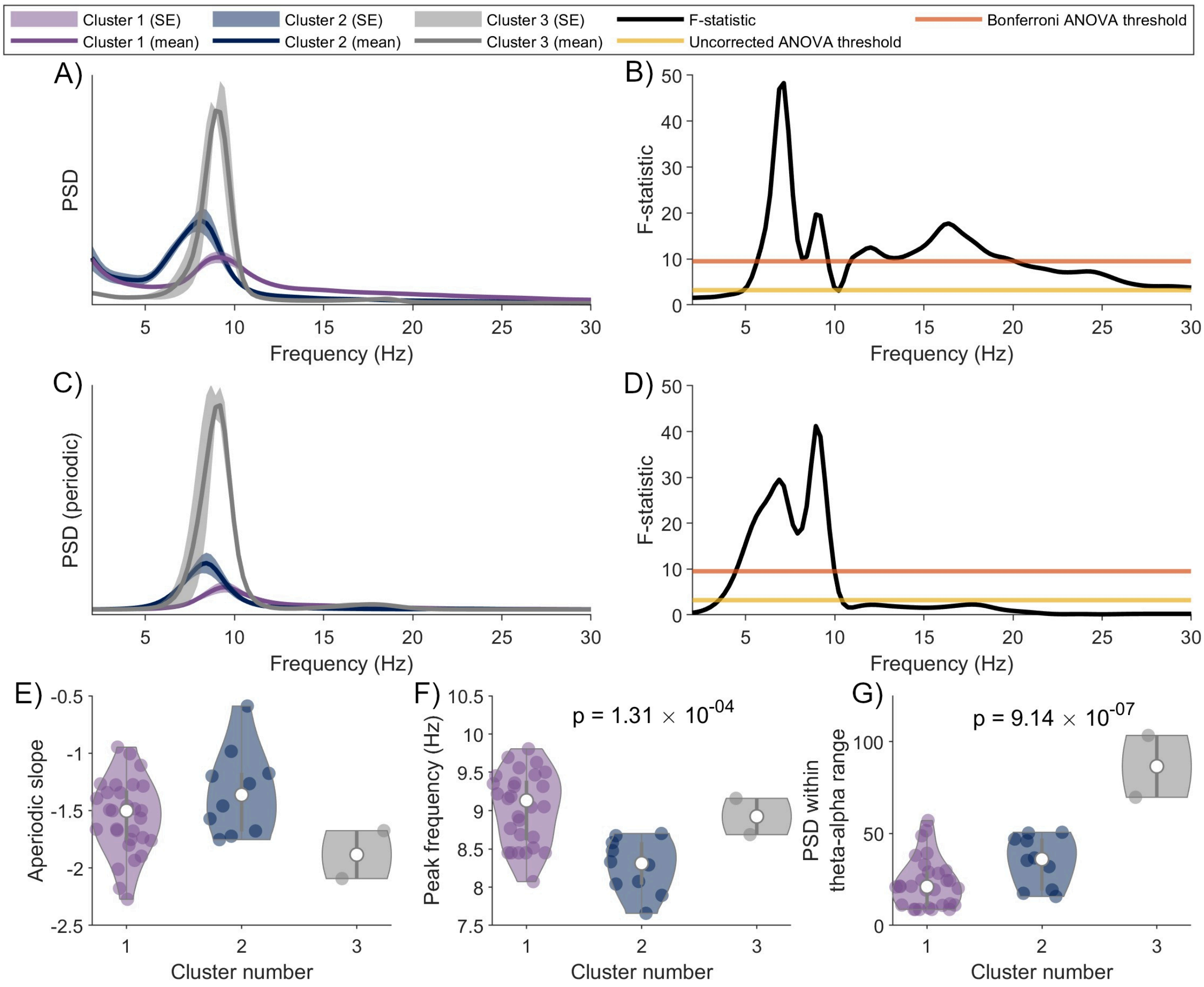


Supplemental Figure S10: **Differences in PSD between clusters are primarily due to theta-alpha frequency oscillations.** A) Mean ($\pm$ standard error) PSD for each of the three spectral clusters. Cluster 1 is characterised by a dominant rhythm around 9 Hz, whereas cluster 2 is characterised by a slowing and small power increase of the oscillation, and cluster 3 is characterised by a large increase in oscillatory power. B) F-statistic for the one-way ANOVA of PSD vs cluster at each frequency shows a broadband difference, with peaks in high and low alpha bands and the beta band. Since broadband changes may be due to relative power normalisation or aperiodic slope, we used spectral parameterisation to separate periodic oscillations and aperiodic 1/f components of the spectrum. C-D) repeat A-B) but for the periodic oscillatory components of the spectrum only. In this case, differences are no longer broadband but primarily at slower theta-alpha frequencies. E) There was no significant difference in aperiodic slope across clusters. At theta-alpha frequencies, there was a significant effect of cluster on F) peak frequency ($F = 11.4$, $p = 1.31 \times 10^{-4}$) and G) periodic power ($F = 20.5$, $p = 9.14 \times 10^{-7}$).

#### Supplemental 3: Connectivity Analysis

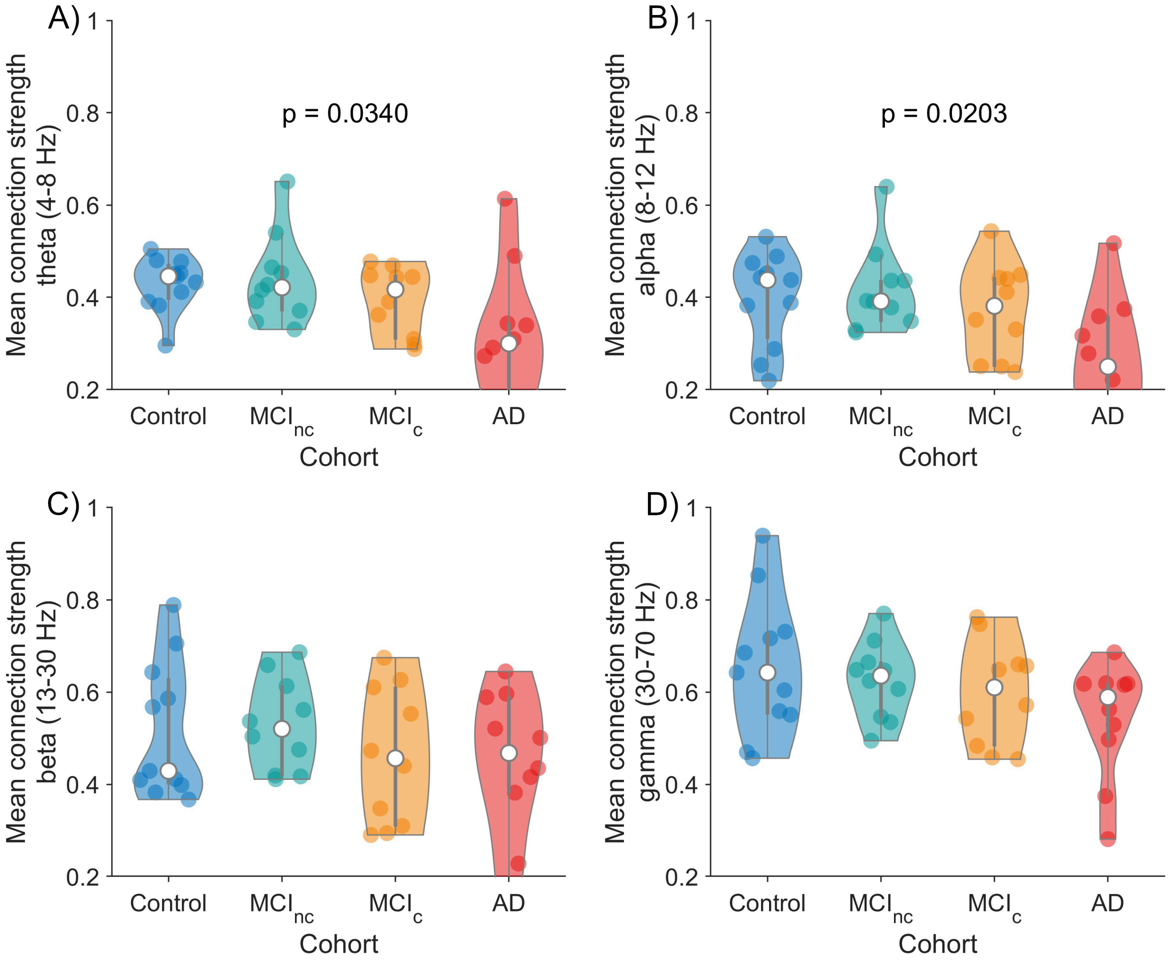


Supplemental Figure S11: **Comparison of mean connectivity across frequency bands for each cohort.** The mean connectivity was compared for each cohort across the A) theta, B) alpha, C) beta and D) gamma frequency bands. Using a one-way ANOVA test, significant differences were observed across the theta ($F = 3.21$, $p = 0.0340$) and alpha ($F = 3.69$, $p = 0.0203$) bands only.

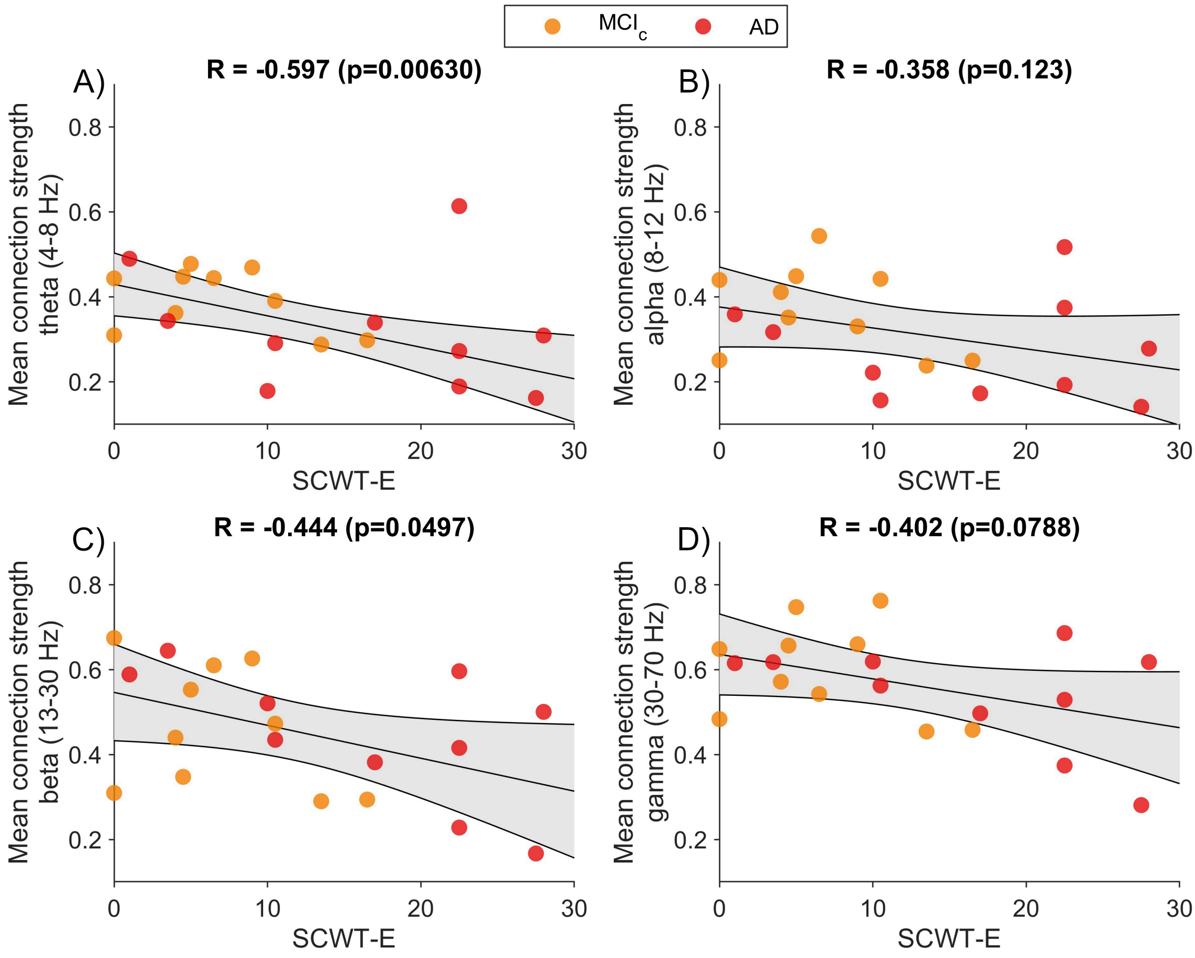


Supplemental Figure S12: **Degree of impairment to SCWT-E scores in people with (prodromal) AD is correlated with functional disconnectivity.** In people with (prodromal) AD (i.e. the AD or MCIC groups), mean connection strength across the 17 nodes of the SCWT attention/working-memory network (see subsection 3.5) significantly negatively correlates with SWCT-E in the A) theta and C) beta bands, but not in the B) alpha and D) gamma bands. Correlations and p-values are shown for each band in the subplot titles. These values were calculated using robust linear regression.

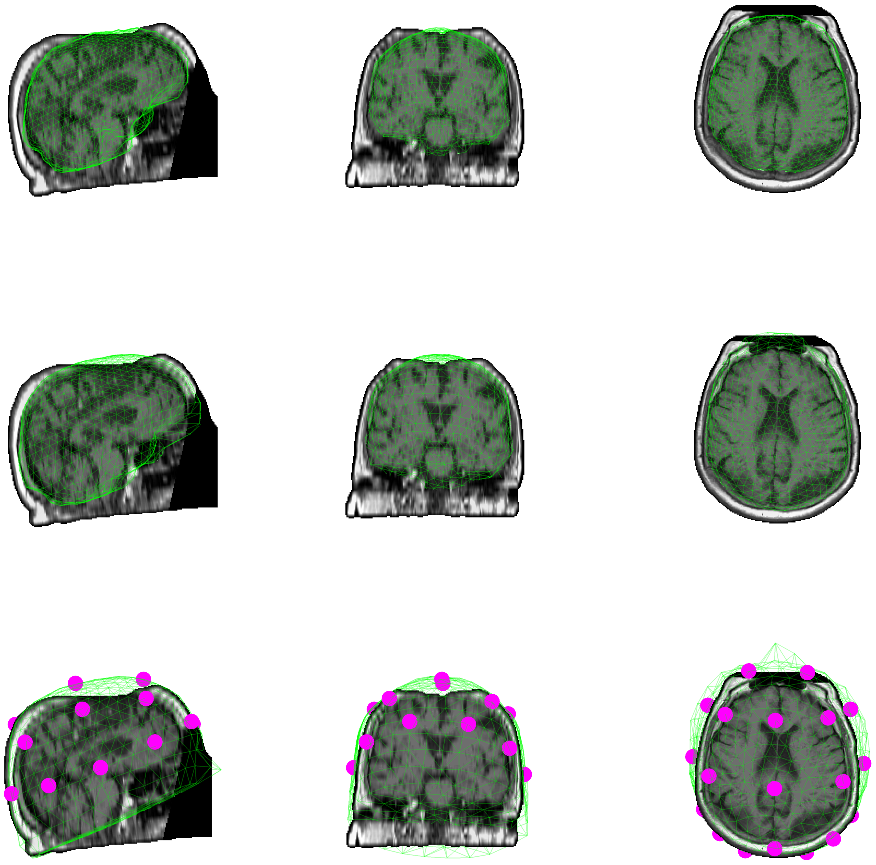

Supplemental Figure S13: **Example 3-layer boundary element method headmodel.** For an example participant (ID number 1), the warped template brain (top row), skull (middle row) and scalp (bottom row) surfaces are shown in green, overlaid on the participant's MRI. Warped template electrode locations are shown in magenta. The MRI is displayed following Freesurfer preprocessing (see Methods). For anonymisation purposes, the MRI has been defacing using SPM12 for plotting.

| Region name | Network | x (mm) | y (mm) | z (mm) | Reference |
|---|---|---|---|---|---|
| l-BA40 | OA | -30 | -62 | 42 | Basten et al. (2011) |
| r-BA40 | OA | 36 | -46 | 42 | Basten et al. (2011) |
| l-FEF/BA8 | OA | -25 | -11 | 54 | Schmidt et al. (2013) |
| r-FEF/BA8 | OA | 27 | -11 | 54 | Schmidt et al. (2013) |
| l-dlPFC | EA | -42 | 6 | 30 | Basten et al. (2011) |
| r-dlPFC | EA | 42 | 6 | 26 | Basten et al. (2011) |
| l-MTG | EA | -62 | -18 | -20 | Conwell et al. (2018) |
| r-MTG | EA | 60 | -4 | -24 | Conwell et al. (2018) |
| SMA | EA | 2 | 1 | 50 | Tzourio-Mazoyer et al. (2002)* |
| l-FIC | EA | -32 | 20 | 0 | Conwell et al. (2018) |
| r-FIC | EA | 32 | 20 | 0 | Conwell et al. (2018) |
| dACC/BA24 | DMN & EA | 2 | 16 | 48 | Basten et al. (2011) |
| l-ParaHippocampal | DMN | -22 | -14 | -24 | Conwell et al. (2018) |
| r-ParaHippocampal | DMN | 22 | -10 | -22 | Conwell et al. (2018) |
| PCC/BA23/BA31 | DMN | -2 | -50 | 25 | Schmidt et al. (2013) |
| l-Angular/BA39 | DMN | -50 | -66 | 28 | Conwell et al. (2018) |
| r-Angular/BA39 | DMN | 50 | -62 | 34 | Conwell et al. (2018) |

Table S2: **MNI coordinates used for nodes in the Stroop network connectivity analysis**. References given are for the choice of MNI coordinates, while the references for choice of regions and association with Stroop scores are given in main text. Network abbreviations: OA=Orienting attention, EA=Executive attention, DMN=Default mode network. Region abbreviations: BA=Brodmann Area, FEF=Frontal Eye Field, dlPFC=Dorsolateral prefrontal cortex, MTG=Medial temporal gyrus, SMA=Supplementary Motor Area, FIC=Frontoinsular cortex, dACC/PCC=dorsal Anterior/Posterior Cingulate Cortex. Prefixes l- and r- refer to left- and right-hemispheres respectively, while those without a prefix are along the medial line of the brain. *SMA MNI coordinates were taken from the centroid of the SMA region in the AAL atlas implemented in Fieldtrip.

#### Supplemental 4: Neural mass modelling of EEG

Neural mass models (NMMs) represent the mean macroscopic behaviour of populations of neurons. In this work, we employ the NMM developed by D T J Liley and Dafilis (2002). This model consists of interacting excitatory and inhibitory neuronal populations and has been used extensively to model EEG rhythms, including the human alpha rhythm (Hartoyo et al., 2019). The model dynamics are governed by differential equations that describe the change in activity (membrane potentials and synaptic dynamics) over time. Each population is considered to be connected to both itself and the other population, with the assumption of fast-acting synapses. Subsequently, for $t \in [0, T]$ with $T \in \mathbb{R}_{>0}$, the model is comprised of the following set of coupled first and second-order differential equations:

$$\tau_e \frac{dh_e(t)}{dt} = h_e^{rest} - h_e(t) + \frac{h_e^{eq} - h_e(t)}{\mid h_e^{eq} - h_e^{rest} \mid} I_{ee}(t) + \frac{h_i^{eq} - h_e(t)}{\mid h_i^{eq} - h_e^{rest} \mid} I_{ie}(t), \tag{2}$$

$$\tau_i \frac{dh_i(t)}{dt} = h_i^{rest} - h_i(t) + \frac{h_e^{eq} - h_i(t)}{\mid h_e^{eq} - h_i^{rest} \mid} I_{ei}(t) + \frac{h_i^{eq} - h_i(t)}{\mid h_i^{eq} - h_i^{rest} \mid} I_{ii}(t), \tag{3}$$

$$\frac{d^2 I_{ee}(t)}{dt^2} + 2\gamma_{ee} \frac{dI_{ee}(t)}{dt} + \gamma_{ee}^2 I_{ee}(t) = \Gamma_e \gamma_{ee} \exp(1)(N_{ee}^{\beta} S_e(h_e(t)) + p(t)), \tag{4}$$

$$\frac{d^2 I_{ei}(t)}{dt^2} + 2\gamma_{ei} \frac{dI_{ei}(t)}{dt} + \gamma_{ei}^2 I_{ei}(t) = \Gamma_e \gamma_{ei} \exp(1)(N_{ei}^{\beta} S_e(h_e(t)) + p_{ei}), \tag{5}$$

$$\frac{d^2 I_{ie}(t)}{dt^2} + 2\gamma_{ie} \frac{dI_{ie}(t)}{dt} + \gamma_{ie}^2 I_{ie}(t) = \Gamma_i \gamma_{ie} \exp(1)(N_{ie}^{\beta} S_i(h_i(t))), \tag{6}$$

$$\frac{d^2 I_{ii}(t)}{dt^2} + 2\gamma_{ii} \frac{dI_{ii}(t)}{dt} + \gamma_{ii}^2 I_{ii}(t) = \Gamma_i \gamma_{ii} \exp(1)(N_{ii}^{\beta} S_i(h_i(t))), \tag{7}$$

where $S(\cdot)$ is a nonlinear sigmoid function that provides a transformation of the mean postsynaptic membrane potential into a mean firing rate. This mapping is defined as,

$$S_j(h_j(t)) = \frac{S_j^{max}}{(1 + exp(-\sqrt{2}\frac{(h_j(t) - \mu_j)}{\sigma_j}))}, \tag{8}$$

for subscripts $j = e, i$ representing the excitatory and inhibitory populations, respectively. The model is numerically solved using the Euler–Maruyama method, in the same way we have described in previous work (Dunstan et al., 2023). Supplemental Table S3 provides a list of all the parameters in the model, along with a physiological interpretation, typically used values and parameter bounds (D T J Liley and Dafilis, 2002; Bojak and Liley, 2005).

All model parameters were optimised by fitting the dynamics of the model to the EEG data. In particular, we followed the approach described in Dunstan et al. (2023), using a multi-objective evolutionary algorithm (MOEA) to fit to two objectives: (1) the sum of squares error in the PSD between model and data; (2) the Kolmogorov–Smirnov test statistic between the data and model node degree of the horizontal visibility graph. We note that to focus on the dominant alpha rhythm, the PSD objective was compared across a frequency range of 5 and 20 Hz. Hyperparameters for the optimization (including population size and generation number) were set to ensure convergence of the algorithm, as detailed in Dunstan et al. (2023).

| Parameter | Interpretation | Typical Value | Range |
|---|---|---|---|
| $h_e^{rest}, h_i^{rest}$ | Mean excitatory/inhibitory membrane potential at rest | $-70$ mV, $-70$ mV | $[-80$ mV,$-60$ mV$]$, $[-80$ mV,$-60$ mV$]$ |
| $N_{ee}^{\beta}$, $N_{ei}^{\beta}$, $N_{ie}^{\beta}$, $N_{ii}^{\beta}$ | Number of excitatory - excitatory/excitatory - inhibitory/inhibitory - excitatory/inhibitory - inhibitory synaptic connections | $4000, 3034, 536, 536$ | $[2000, 5000]$, $[2000, 5000]$, $[100, 1000]$, $[100, 1000]$ |
| $\Gamma_e, \Gamma_i$ | Excitatory/inhibitory mean synaptic gain | $0.4$ mV, $0.8$ mV | $[0.1$ mV, $2$ mV$]$, $[0.1$ mV, $2$ mV$]$ |
| $\gamma_e, \gamma_i$ | Excitatory/inhibitory postsynaptic potential rate constant | $0.3$ /ms, $0.065$ /ms | $[0.1$ /ms,1 /ms$]$, $[0.01$ /ms, $0.5$ /ms$]$ |
| $\tau_e, \tau_i$ | Passive excitatory/inhibitory membrane time constant | $10$ ms, $10$ ms | $[5$ ms, $150$ ms$]$, $[5$ ms, $150$ ms$]$ |
| $S_e^{max}$, $S_i^{max}$ | Maximum mean firing rate of excitatory/inhibitory population | $0.5$ /ms, $0.5$ /ms | $[0.05$ /ms, $0.5$ /ms$]$, $[0.05$ /ms, $0.5$ /ms$]$ |
| $\mu_e, \mu_i$ | Excitatory/inhibitory firing rate thresholds | $-50$ mV, $-50$ mV | $[-55$ mV, $-40$ mV$]$, $[-55$ mV, $-40$ mV$]$ |
| $\sigma_e, \sigma_i$ | Excitatory/inhibitory firing threshold standard deviation | $5$ mV, $5$ mV | $[2$ mV, $7$ mV$]$, $[2$ mV, $7$ mV$]$ |
| $h_e^{eq}, h_i^{eq}$ | Excitatory/inhibitory mean reversal potential | - | $[-20$ mV, $10$ mV$]$, $[-90$ mV, $-65$ mV$]$ |
| $p_{ee}, p_{ei}$ | Rate of excitatory-excitatory/excitatory-inhibitory input | - | $[0$ /ms, $10$ /ms$]$, $[0$ /ms, $10$ /ms$]$ |
| $\xi$ | Standard deviation of noise perturbation | - | $[0, 10]$ |

Table S3: **Parameters in the Liley neural mass model.** The parameter ranges (as defined in D T J Liley and Dafilis (2002); Bojak and Liley (2005)) were those used in the MOEA.

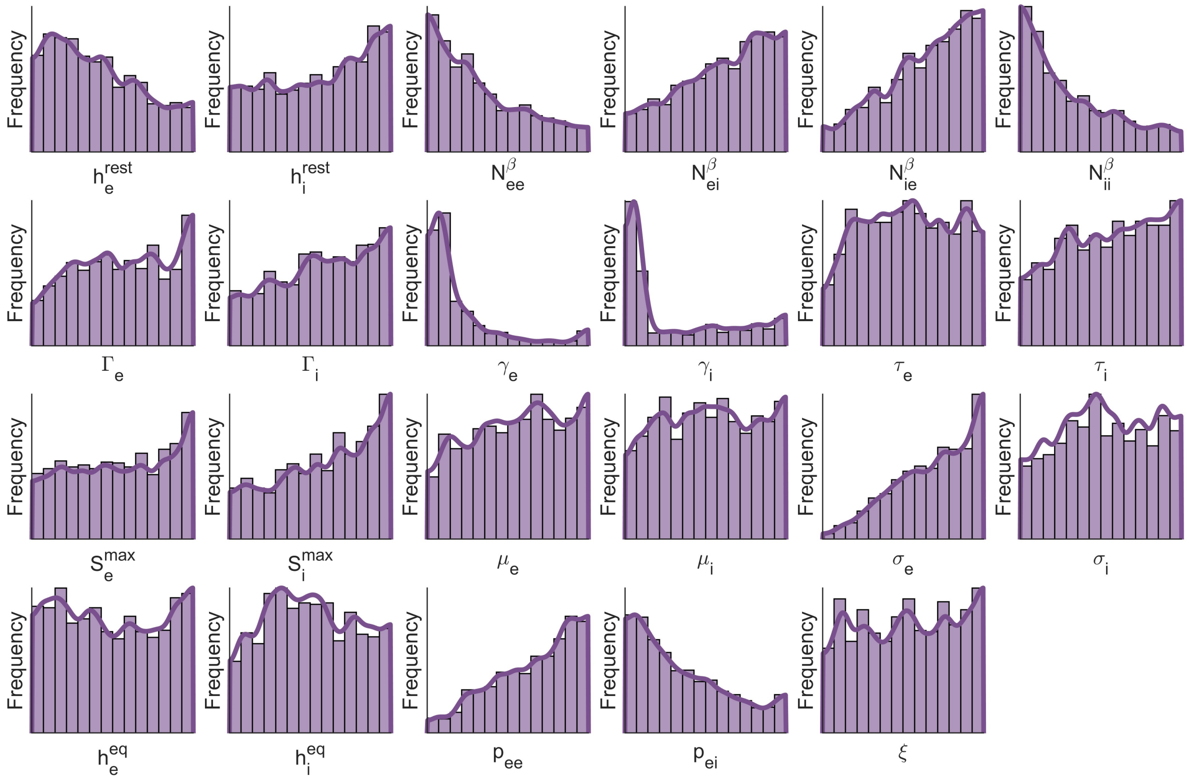


Supplemental Figure S14: **Parameters recovered from fitting to cluster 1.** Distributions show parameters recovered using the MOEA to fit the Liley model to all EEG data in cluster 1.

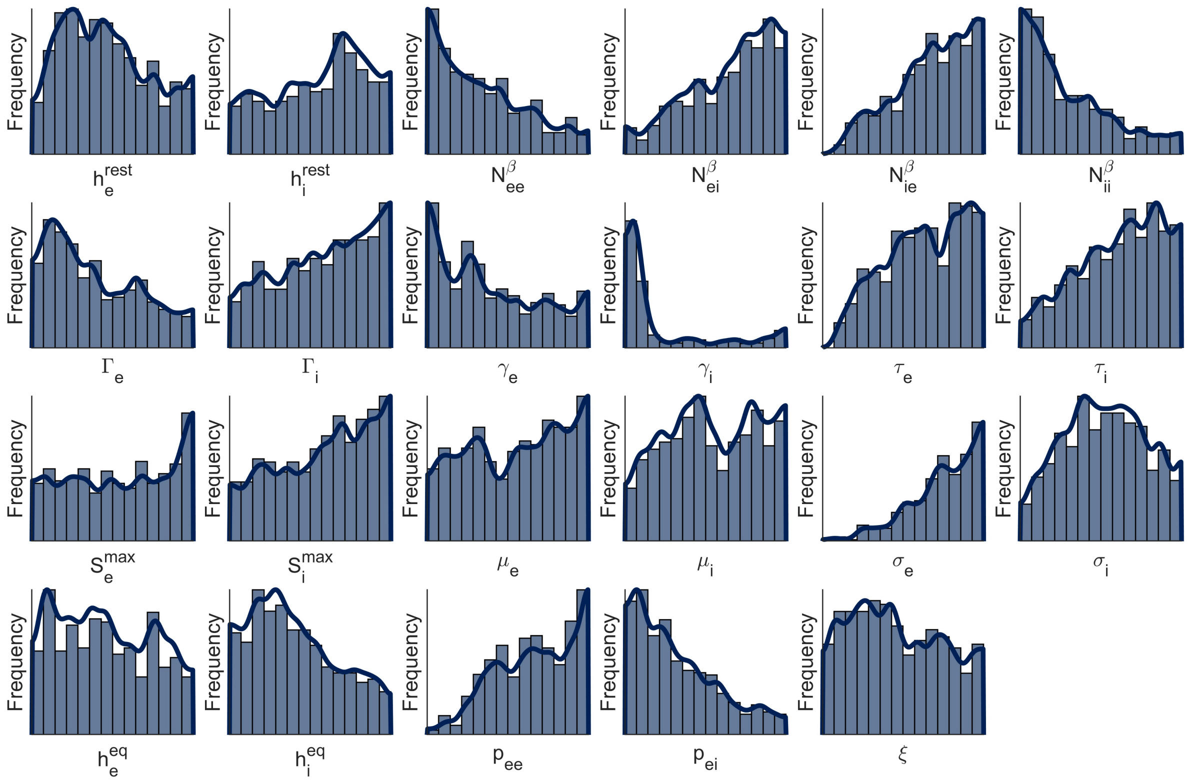


Supplemental Figure S15: **Parameters recovered from fitting to cluster 2.** Distributions show parameters recovered using the MOEA to fit the Liley model to all EEG data in cluster 2.

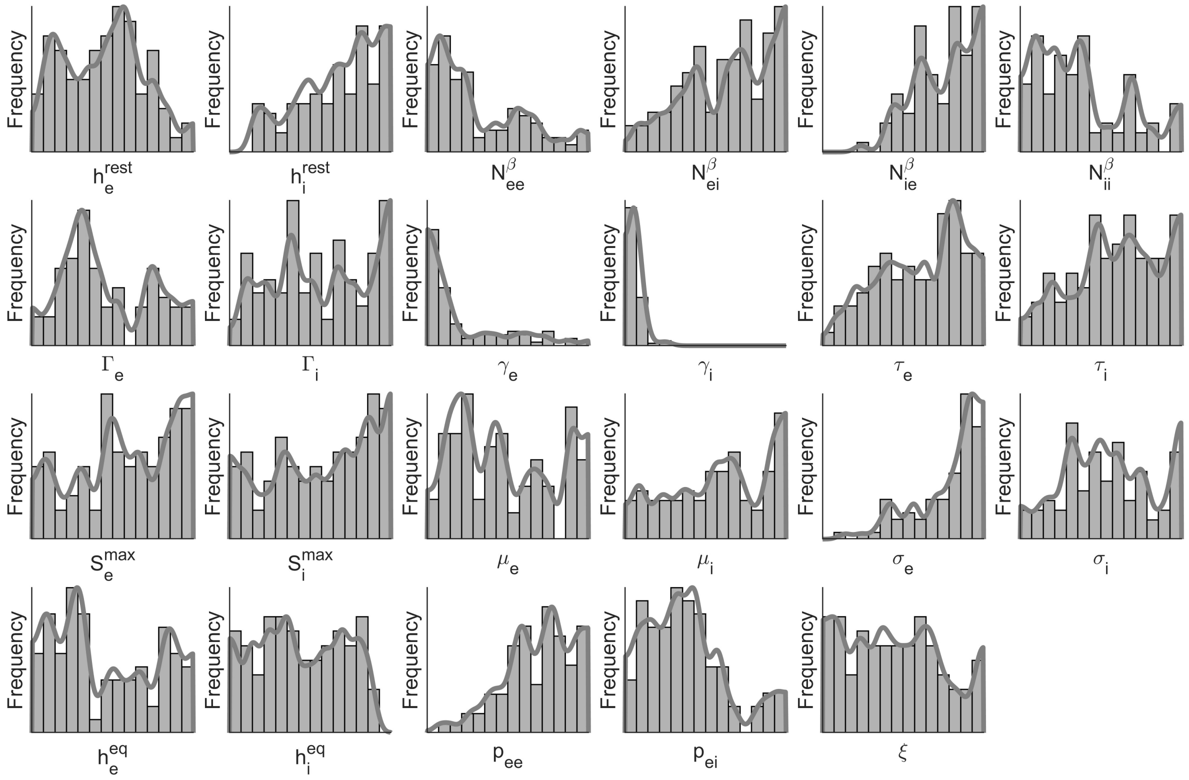


Supplemental Figure S16: **Parameters recovered from fitting to cluster 3.** Distributions show parameters recovered using the MOEA to fit the Liley model to all EEG data in cluster 3.
